# COMMON AND DISTINCT NEURAL CORRELATES OF SOCIAL INTERACTION PERCEPTION AND THEORY OF MIND

**DOI:** 10.1101/2024.12.19.628993

**Authors:** Zizhuang Miao, Heejung Jung, Philip A. Kragel, Ke Bo, Patrick Sadil, Martin A. Lindquist, Tor D. Wager

## Abstract

Social interaction perception and theory of mind (ToM) frequently co-occur, but their commonalities and distinctions at behavioral and neural levels remain unclear. Participants (*N* = 231) provided moment-by-moment ratings of four text and four audio narratives on social interactions and ToM engagement, which were reliable (split-half *r* = .98 and .92, respectively) but only modestly correlated (*r* = .32). In a second sample (*N* = 90), we analyzed the co-variation between social interaction and ToM ratings and fMRI activity during text and audio narratives. Social interaction and ToM activity maps generalized across modalities (spatial *r* = .83 and .57, respectively), both with significant, overlapping clusters in canonical mentalizing regions (FDR *q* < .01). ToM uniquely engaged the lateral occipitotemporal cortex, left anterior intraparietal sulcus, and right premotor cortex. These results suggest that perceiving social interactions automatically involves mentalizing, and ToM additionally engages brain regions for action understanding and executive functions.

## Common and distinct neural correlates of social interaction perception and theory of mind

As social animals, we are surrounded by others whose appearances, expressions, and actions we can perceive, and whose thoughts, beliefs, and emotions we can infer. Based on a processing-stage theoretical framework (or dual-process model; see Lieberman, 2007; Bohl & van den Bos, 2012; Lieberman, 2022), there are two types of social cognitive processes: early perceptual processes that are automatic, inflexible, and specific to sensory modality, and late inferential processes that are controlled, flexible, and operate on multi-modal information. Although qualitatively different, the two processes often happen simultaneously and transition seamlessly. For example, people can readily infer animacy and mental states while perceiving the motions and interactions of completely inanimate geometric shapes (Heider & Simmel, 1944). However, social cognition research has been undergoing a separation, with some studies focusing on automatic social perception (e.g., Pitcher & Ungerleider, 2021) and others on controlled social inferences (e.g., Schurz et al., 2021). One prominent example is the distinction between social interaction perception, which involves perceiving the interactions of other agents from a third-person perspective (e.g., Isik et al., 2017), and theory of mind (ToM), which involves deliberating over other agents’ mental states (e.g., Frith & Frith, 2005). The separation in research topics has promoted the idea that social interaction perception and ToM are distinct mental processes implemented in separate brain regions (McMahon & Isik, 2023). Given their frequent co-occurrence (Heider & Simmel, 1944) and potential confounding (e.g., Varrier & Finn (2022) and Moessnang et al. (2016) studied social interaction perception and ToM, respectively, using similar stimuli), results from different studies may not reflect their actual differences. It thus remains unclear how common or unique social interaction perception and ToM are at both neural and behavioral levels.

We will divide this question into two parts and seek answers by modeling social interaction perception and ToM in the same neuroimaging study. First, we will investigate to what extent social interaction perception is a purely perceptual process or also involves conceptual processes. Social interaction perception has been predominantly studied using visual or audiovisual stimuli, such as animations of geometric shapes (e.g., Varrier & Finn, 2022), animations of virtual characters (e.g., Schilbach et al., 2006), videotapes (e.g., Ainsworth et al., 2021), and movie clips (e.g., Lahnakoski et al., 2012). It activates many heavily visual regions, such as middle temporal cortex or lateral occipitotemporal cortex (e.g., Landsiedel et al., 2022; Wurm & Caramazza, 2019) and cuneus (e.g., Schilbach et al., 2006; Wolf et al., 2018). Accumulating behavioral and neural evidence suggests that social interaction perception is a visually grounded perceptual process (McMahon & Isik, 2023). However, other neuroimaging studies have opposite findings: brain regions that respond to social interactions overlap with those responding to ToM (Arioli & Canessa, 2019), include multimodal regions in anterior temporal cortex (McMahon et al., 2023), and show generalizability across sensory modality (Landsiedel & Koldewyn, 2023) and task modality (Lee Masson et al., 2024b). These findings implicate an alternative view that social interaction perception has cross-modality components and could involve conceptual processing. To further compare these two views, we investigate whether any brain regions that respond to social interactions in visual stimuli in previous studies also respond to visually and auditorily presented verbal stimuli in the current study. We also compare the neural correlates of social interaction perception with those of ToM which, by definition, involves conceptual inferences.

Second, we will ask to what extent ToM is a controlled or automatic process. Much developmental and evolutionary evidence (e.g., Frith & Frith, 2005; Richardson et al., 2018; Deen et al., 2023) has shown that ToM is a more resource-demanding mental function compared to more primitive functions such as face perception. Neuroimaging evidence supports this by showing that ToM is primarily implemented in the multimodal neocortex, such as temporoparietal junction, medial prefrontal cortex, and precuneus (e.g., Schurz et al., 2021; Dufour et al., 2013). Based on this view, existing studies have operationalized ToM as a controlled mental process, such as deliberately reasoning about the beliefs of characters in stories (Saxe & Kanwisher, 2003) and the intentions of people in pictures (Spunt & Adolphs, 2014). However, when researchers do not ask participants to deliberately use ToM while watching a movie, the ToM annotations correlate with neural activity in largely the same brain regions found in controlled experiments (Jacoby et al., 2016). These regions are also engaged by many other pre-reflective subjective construal processes (Lieberman, 2022). The findings suggest that ToM may be used in an effortless, automatic way during naturalistic experience. To better understand how controlled or automatic ToM is, we study it in situations akin to real life using narratives and compare it with another more automatic process, social interaction perception.

Addressing similar questions, several recent studies have made breakthroughs by directly comparing the neural correlates of social interaction perception and ToM and found that they are separable (Isik et al., 2017; Lee Masson et al., 2021; Lee Masson et al., 2024a). This suggests that social interaction perception and ToM are at different processing stages, as separation in neural implementations appears to be a sufficient condition for separation at the mental level. However, a methodological limitation is worth noting: previous studies have operationalized ToM in controlled experiments as one condition versus another (Isik et al., 2017), and in movie stimuli as the annotations of where the viewer is led to think about the characters’ thoughts (Jacoby et al., 2016) or where the characters are inferring other characters’ mental states (Lee Masson & Isik, 2021; Lee Masson et al., 2024a). These operational definitions are more like a stimulus property, or “ToM demands,” rather than the underlying cognitive process, or ToM *per se*. This discrepancy between measurements and the mental process could impact the validity and sensitivity of linking ToM to neural activity (Zaki & Ochsner, 2009). Indeed, in the same dataset, voxel-wise modeling reveals separable neural correlates between ToM and social interaction perception (Lee Masson & Isik, 2021), but regions-of-interest-based analysis finds some overlaps (Lee Masson et al., 2024a), partly because the latter method has larger power. To overcome this limitation, we use self-report measurement for ToM during narratives and compare it with the annotation measurement used in previous studies.

Taken together, we aim to investigate to what extent social interaction perception is a modality-specific perceptual process and ToM is a controlled process. For this, we took a “cross-modal replication strategy” by presenting written narratives to participants as both texts and audios, and compared neural correlates across the two modalities. We also sought to identify specific regions for social interaction perception and ToM regardless of sensory inputs. We correlated social interactions and ToM with neural activity in a single model and compared their neural correlates when controlling for each other. Spatially separable neural correlates of social interaction perception and ToM can support the hypothesis that social interaction perception and ToM are two distinct mental processes, whereas shared neural correlates between the two will challenge it.

## Results

We collected and analyzed two datasets in the current study. The first dataset came from an online behavioral experiment where participants (*N* = 231) provided ratings about how much they thought there were social interactions (*N* = 114) or how much they were using ToM (*N* = 117) in real time as they read four written narratives and listened to another four. We extracted the medians of those ratings to serve as continuous estimates of the subjective perception of social interactions and engagement of ToM for general populations. The second dataset came from a neuroimaging experiment where another group of participants (*N* = 90) passively experienced the same narratives under functional magnetic resonance imaging (fMRI) scanning without providing any ratings. We then correlated the medians of online behavioral ratings with fMRI data using voxel-wise random-effect general linear models (GLM; Figure 1a). All inferential statistical tests were two-tailed. Significance in multiple comparisons in voxel-wise modeling of neural data was corrected by false discovery rate (FDR; Benjamini & Hochberg, 1995) *q* < .01. Significance in other multiple comparisons was corrected using Bonferroni correction.

**Figure 1.**
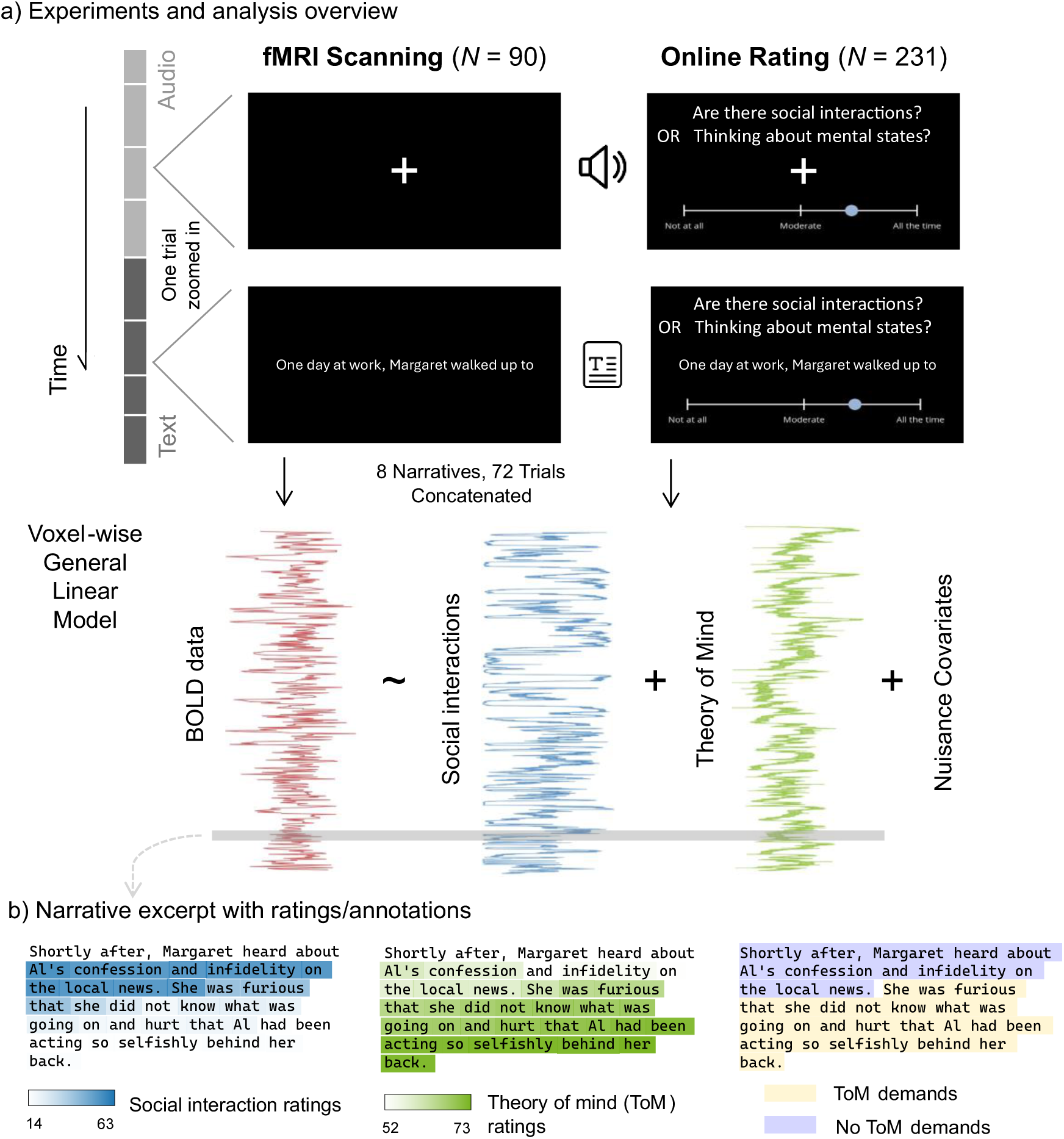
Study overview. a) The stimuli used in this study were eight narratives, with the first four presented as audios and the last four presented as texts. One group of participants (*N* = 90) experienced the narratives during functional magnetic resonance imaging (fMRI) scan. Another group of participants (*N* = 231) experienced the narratives online on their personal computers and simultaneously provided moment-by-moment ratings about social interactions in the narratives (*N* = 114) or their use of theory of mind (ToM) (*N* = 117). The black rectangles illustrate the screen that participants saw when narratives were presented. One online participant only saw one of the two questions shown in this figure throughout the study. We concatenated blood-oxygen-level-dependent (BOLD) signals in fMRI scanning (one fake BOLD time series was plotted for illustration only) and the medians of online ratings across all trials (actual data were plotted). Then we located the neural correlates of social interaction perception and ToM by two-stage voxel-wise general linear models. Shaded areas around the lines of online ratings indicate confidence intervals calculated by median absolute difference. b) An example narrative excerpt extracted from the eighth trial of Narrative #1 (see Supplementary Materials for all narrative texts). The medians of social interaction and ToM ratings during this e cerpt, as well as researchers’ annotations of ToM demands, were displayed as the highlight color of each word. The approximate time window of this excerpt was shown by the horizontal gray bar in Panel a.

In the following sections, we first validate the methods by checking whether written narratives and self-report measurement could reliably reveal neural correlates of social interaction perception and ToM. Then we directly compare the neural correlates of social interaction perception and ToM and use Bayes Factors to classify each voxel as responding to both social interactions and ToM or only to one of them. We found that the neural correlates of social interaction perception were highly similar across modalities. Furthermore, the voxels that responded to social interactions across modalities were also associated with ToM, while ToM was uniquely associated with other brain regions.

### Cross-modal generalizability of social interaction perception

Because few existing studies have presented social interactions in written narratives, we tested whether the narratives were valid and sensitive to probe the neural correlates of perceiving social interactions.

#### Subheader 1. Consistent and reliable ratings of social interactions

In the online study, participants’ ratings were similar to each other (Figure 2a left). This was corroborated by the moderate to high pairwise correlations between participants’ ratings (Median = .47). Participants most frequently used the two ends of the scale to answer the social interaction question (Figure 2a right), suggesting that they tended to have high confidence about the presence or absence of social interactions. More importantly, participants provided highly reliable ratings, as suggested by large split-half correlations ([.96, .99], *M* = .98; estimated from 2000 permutation samples and corrected by the Spearman-Brown formula (Brown, 1910; Spearman, 1910)). Overall, these results suggest that participants perceive social interactions in the written narratives in a consistent and reliable way.

**Figure 2.**
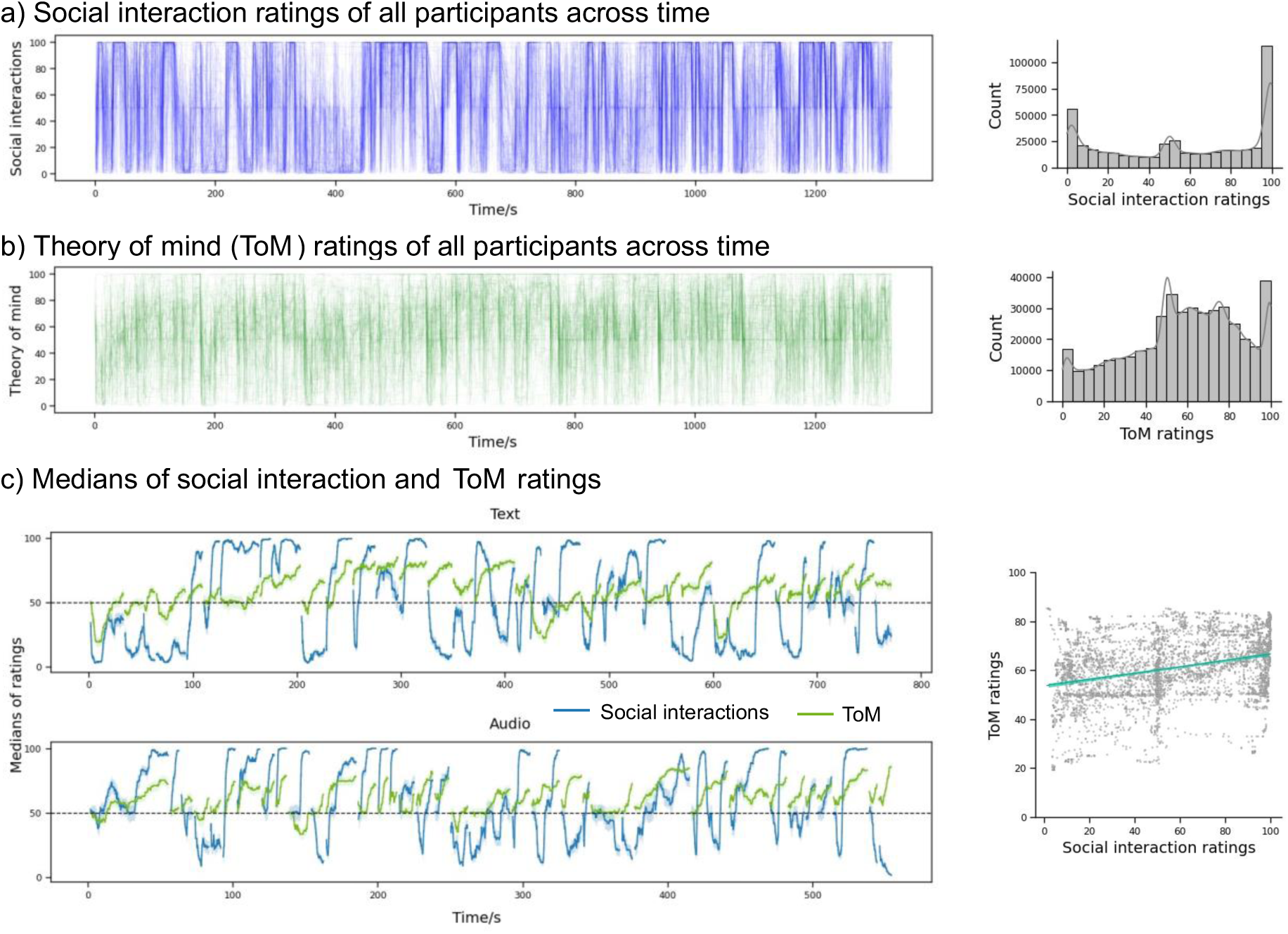
Ratings of online participants about social interactions and theory of mind (ToM). and b) In the line plots on the left, the moment-by-moment ratings of one participant were plotted as a thin quasi-transparent line. Thicker/darker parts indicate that more participants shared ratings at those time points. In the histograms on the right, all ratings were binned and plotted. c) In the line plots on the left, medians of ratings were shown by solid lines, and confidence intervals of the medians were shown by shaded areas around the line. We divided all ratings into two modalities and showed them in two plots. Note: the lines have many discontinuity points because a few data points at the start of each trial were removed from further analysis due to inadequate valid ratings (see Methods for details) and we show the version after removal in this figure. In the scatter plot on the right, each dot corresponds to a time point in the line plots. The cyan line is the best fit line estimated by ordinary least squares regression (*r* = .323).

#### Subheader 2. Similar neural correlates of social interaction perception across modalities and with previous findings in narratives

We used medians of online participants’ ratings in the Te t and udio modality separately to regress each fMRI participant’s neural activity. Then we performed group-level t-tests (*N* = 90) over the individual regression coefficients for the two modalities separately. We found that the unthresholded whole-brain effect maps were highly similar across modalities (*r* = .83; for thresholded maps see Figure 3a left). In a side analysis, we also found that the whole-brain predictive models for social interactions were generalizable across modalities (see Supplementary Results). This suggests that participants use the same neural circuitries to process social interaction information both when reading and when listening to the narratives.

**Figure 3.**
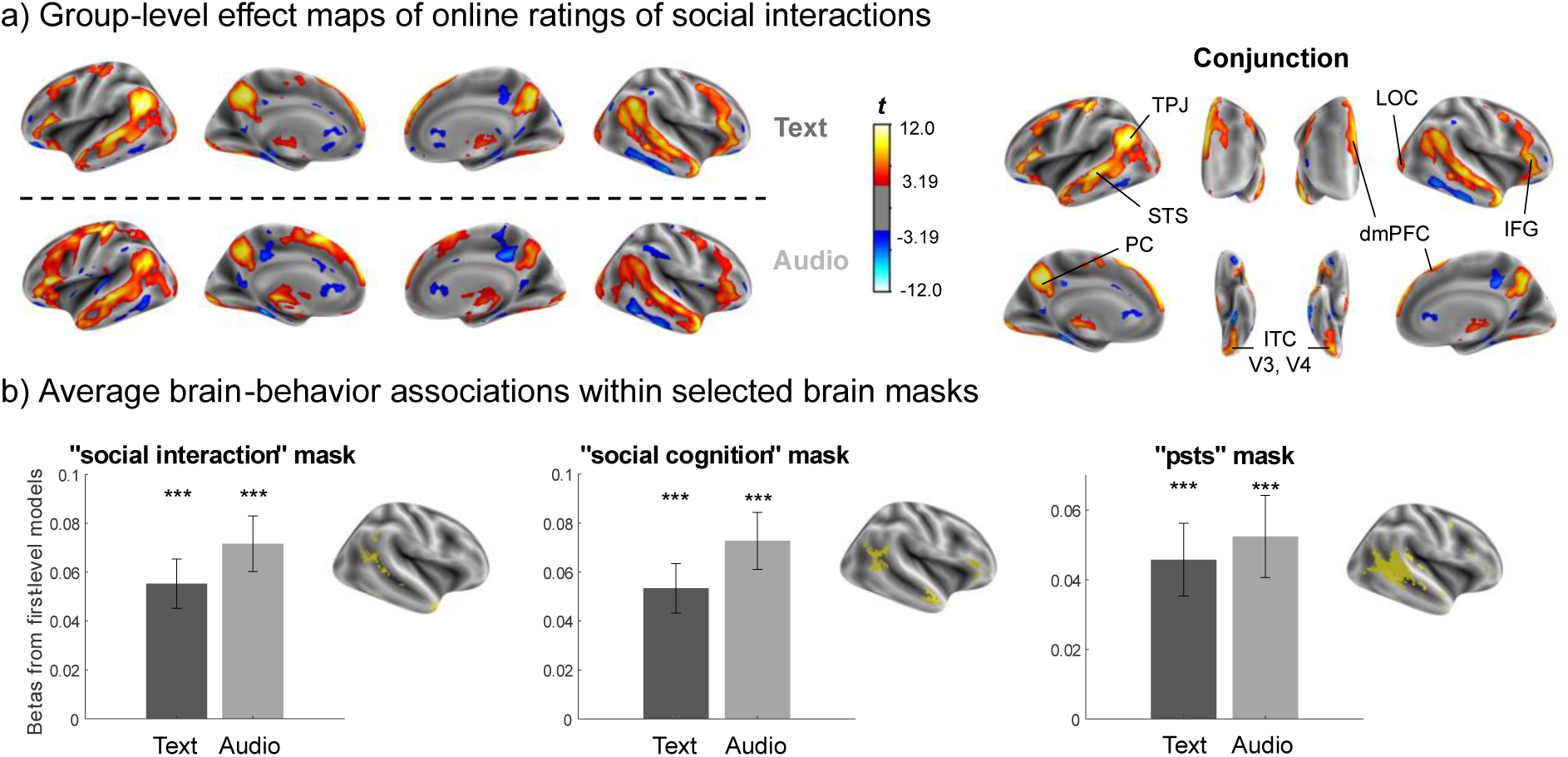
Neural correlates of social interaction perception in narratives. a) Group-level effect maps of social interactions as the medians of online participants’ ratings, thresholded at FDR *q* < .01. Maps on the left showed separate patterns for Text and Audio modality, and maps on the right showed the conjunction across modalities. We did analyses in the volumetric space and projected results onto the Freesurfer average surface maps (Fischl, 2012) by linear interpolation for visualization only. The colormap of *t* values applied to all maps. We annotated several key regions that were significant in both hemispheres and both modalities. Abbreviations: TPJ – temporoparietal junction; STS – superior temporal sulcus; PC – precuneus; dmPFC – dorsomedial prefrontal cortex; IFG – inferior frontal gyrus; LOC – lateral occipital cortex; ITC – inferior temporal cortex. b) Group-level effects of social interactions in two modalities within three brain masks. Parts of each mask were illustrated by the yellowish-green fill color in the surface plots of the right hemisphere; the whole masks were used for analysis. Bar heights indicate the means of regression coefficients (betas) from the first-level models within each mask across participants, which reflect the strength of associations between social interaction ratings and neural activity within each mask. Error bars indicate 95% confidence intervals. *** indicates *p* < .001 in one-sample *t*-tests against 0 (after Bonferroni correction).

We then located the cross-modal neural correlates of social interaction perception by taking the conjunction of the two effect maps (Figure 3a left). In the conjunction map, a voxel was significant if and only if it was significant in both original maps with the same sign. Many regions implicated in social cognition studies were significant in the conjunction map, including bilateral temporoparietal junction (TPJ), superior temporal sulcus (STS), precuneus (PC), dorsomedial prefrontal cortex (dmPFC), and inferior frontal gyrus (IFG). Moreover, early visual cortex including V3 and V4, lateral occipital cortex (LOC), and parts of ventral visual stream or inferior temporal cortex (ITC) (Glasser et al., 2016) were also significant (Figure 3a right, also see Table S1). This suggests that social interactions in narratives are correlated with neural activity in regions typically seen as responsible for visual processing, even when there is no visual information about social interactions.

The cross-modal regions responding to social interactions in the narratives were similar to those reported in previous studies (e.g., Georgescu et al., 2014; Walbrin et al., 2018; Wagner et al., 2016). To further compare the results with previous findings in a quantitative way, we used the Neurosynth database (Yarkoni et al., 2011) to identify brain masks where all vo els were significantly related to “social interaction”, “social cognition”, or “psts” (Isik et al., 201 ; Pitcher & Ungerleider, 2021). Then we e tracted the means of regression coefficients (betas) of social interactions within the masks, a metric reflecting the association between within-mask neural activity and behavioral ratings of social interactions. The brain-behavior associations under both modalities within all three masks were significantly above 0 (all *t*(89)s > 6.5, all *ps* < .001; Figure 3b). This indicates that brain areas associated with social interactions in previous studies are also activated by the social interaction ratings in the current study. We further tested the robustness of the results by annotating social interactions in the narratives following the methods of previous studies (e.g., Nijhof & Willems, 2015; Lee Masson & Isik, 2021) and found similar result patterns (see Supplementary Results). In summary, the results at both behavioral and neural levels support that narratives are valid and sensitive stimuli to probe the neural correlates of social interaction perception.

### Validation of self-report measurement of ToM

In the Introduction, we argued that researchers’ annotations of ToM events should be thought of as “ToM demands” and may not be sufficiently sensitive to locali e ToM-related neural activity. We sought to overcome this limitation by asking participants to report how much they were thinking about characters’ mental states, and refer to this measurement as “ToM engagement”. We also annotated the narratives in terms of ToM demands (Figure 1b) which approximated the annotations in published studies (e.g., Lee Masson & Isik, 2021). Below we quantitatively compare the two measurements, ToM engagement and ToM demands, on the behavioral and neural levels.

#### Subheader 1. Low similarity between ToM engagement and ToM demands

For ToM engagement ratings, participants used the middle to right end portion of the scale more than the social interaction ratings, suggested by denser distributions in the range 50-80 (Figure 2b) and smaller variance across time (Mean standard deviation = 19.35 for ToM engagement; 32.97 for social interactions). Participants were less similar to one another on their ToM engagement ratings, as seen in lower pairwise correlations between participants (Median = .21) than social interactions. Still, the ToM engagement ratings were highly reliable (split-half correlations [.86, .95] in 2000 permutation samples, *M* = .92, corrected by the Spearman-Brown formula), supporting the validity of applying those ratings to another participant group.

To obtain ratings for ToM demands, four researchers in our group independently annotated the narratives in the unit of sentence parts, the smallest chunk in a sentence that contains one or one sequence of predicates and their associated objects (Nijhof & Willems, 2015). We assigned every word in the narrative into one sentence part. The raters determined whether each sentence part had explicitly mentioned mental states of characters (1 yes, 0 no), as the labels for “ToM demands”. In the current report, “mental states” included both the cognitive aspects (e.g., beliefs and goals), and the affective aspects (e.g., emotions and desires), and were typically signaled by specific verbs and adjectives (see Figure 1b for an example). Overall, the raters gave moderately similar annotations about ToM demands (pairwise correlations between raters ranged from .51 to .60 (*M* = .57) and the Fleiss’ kappa was .54). Then we grouped all the annotations and assigned a consensus label to each sentence part (conflicts were addressed by discussion).

We compared the ToM demand labels to the medians of ToM engagement ratings under the Text and Audio modalities separately. Correlations between the two measures were not significant (*r* = .14, *p* = .22 for Text, and *r* = .19, *p* = .09 for Audio; *p* values estimated from 10000 bootstrapped phase randomization samples). This indicates that ToM demands annotations capture different behavioral profiles than participants’ usage of ToM.

#### Subheader 2. Higher sensitivity of ToM engagement to localize ToM-related neural activity

Given that ToM engagement and ToM demands may be qualitatively different on the behavioral level, we expected to find different neural correlates associated with them. The question is which of the two is better at locating neural activity related to ToM. We answered this by comparing the neural correlates of the two measures both qualitatively (checking which regions were significant) and quantitatively (within brain masks associated with ToM in previous studies). Specifically, we fit two GLMs with the only difference being whether the ToM regressors were related to ToM engagement or ToM demands, and we compared neural results across sensory modalities. For the qualitative comparison, we found that ToM engagement was significantly correlated with bilateral TPJ, STS, PC, dmPFC, and IFG in both modalities, the typical regions associated with ToM (Schurz et al., 2021). By contrast, ToM demands only showed cross-modality activations in a subset of those regions, including bilateral PC and STS, left TPJ, and left IFG, and the areas of significant regions were smaller under the same threshold (Figure 4a).

**Figure 4.**
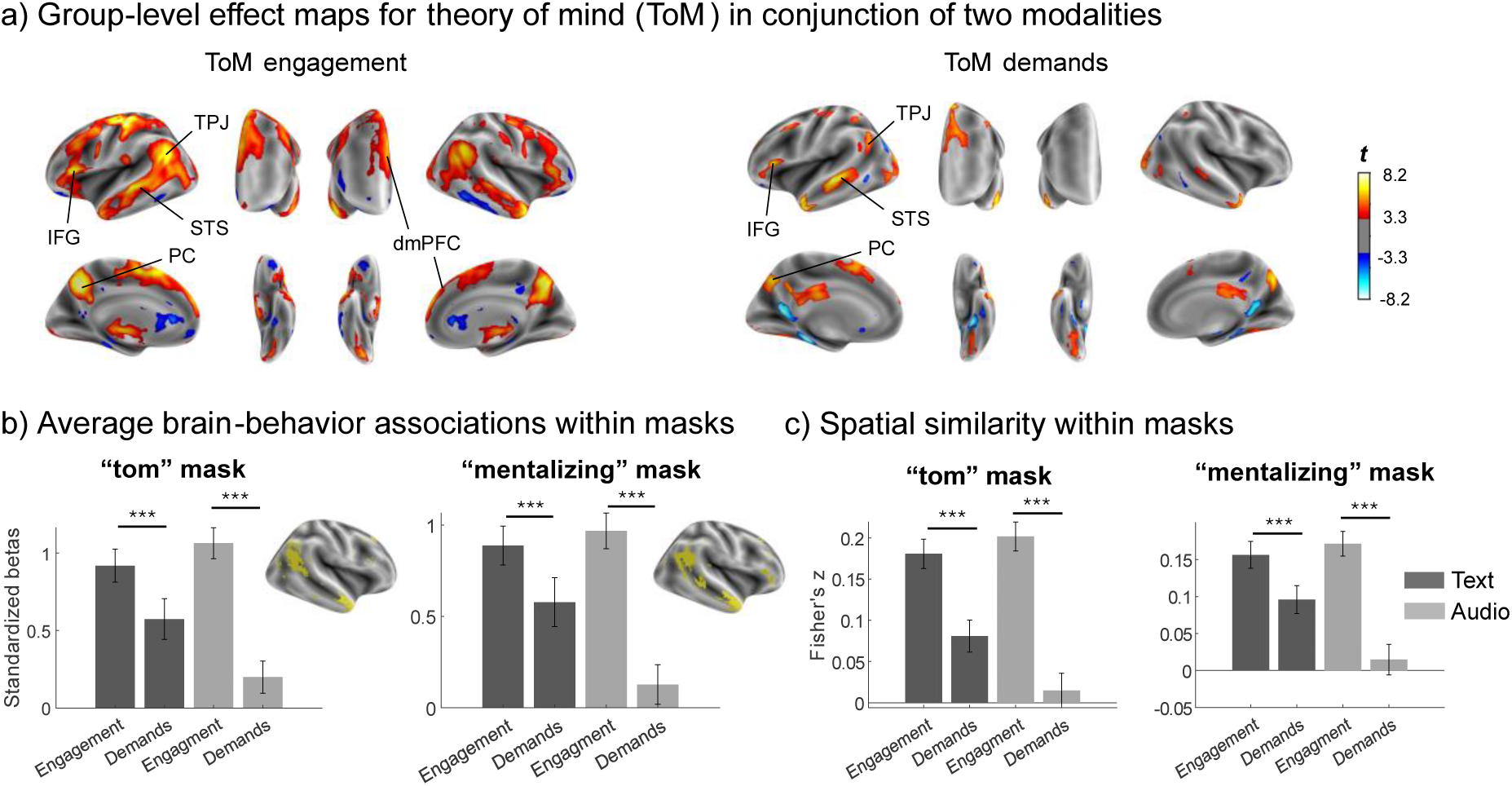
Neural correlates of theory of mind (ToM) engagement and ToM demands. a) Group-level effect maps of ToM as the medians of online participants’ ratings of ToM engagement and the offline annotations of ToM demands, thresholded at FDR *q* < .01. We only displayed the conjunction maps across modalities for simplicity. The method for projection from volumetric space to surface space and abbreviations were the same as in Figure 3a. b) Group-level comparisons of the first-level beta estimates for ToM engagement (“ ngagement”) and ToM demands (“ emands”) in two modalities within two brain masks. Bar heights indicate means of standardized betas and error bars indicate 95% confidence intervals. Parts of the masks on the right hemisphere were indicated by the yellowish-green fill color in the surface plots. c) Group-level spatial pattern correlations between beta maps and Neurosynth maps within each mask. Bar heights indicate the means of Fisher *z* values transformed from Pearson’s correlation coefficients and error bars indicate 95% confidence intervals. In all graphs, *** indicates *p* < .001 (after Bonferroni correction).

For the quantitative comparison, we selected brain masks from the “tom” and “mentali ing” term maps from the eurosynth database (Yarkoni et al., 2011) and the ToM group maps from a false belief task (Dufour et al., 2013). The first comparison was between the brain-behavior associations within masks, operationalized as the average standardized regression coefficients (betas). Standardized betas were calculated as each vo el’s beta divided by the average of the absolute betas in the whole brain, which were at roughly the same scale and comparable across models. The average standardized betas of ToM engagement and ToM demands did not differ in the whole brain (*t*(89) = 1.51, *p* = .27 for Text; *t*(89) = 2.04, *p* = .09 for Audio). By contrast, in almost all selected masks and both modalities, ToM engagement had larger standardized betas than ToM demands (all *t*(89)s > 3, all *p*s < .05), with the only exception in the right STS mask under the Text modality (*t*(89) = 1.34, *p* > .05; results from the Neurosynth masks were shown in Figure 4b, other results were in Figure S4). This indicates that ToM engagement is associated with neural activity in canonical ToM regions more strongly than ToM demands.

The second comparison was on the similarity between the spatial patterns of betas and meta-analytic maps within masks. In each mask created from the Neurosynth database, we e tracted each participant’s betas and correlated them with the *z* values in the meta-analytic map. We then transformed the Pearson’s correlation coefficients to Fisher’s *z* values to ensure normality, which were then subject to paired-sample t-tests. In both “tom” and “mentali ing” masks and both modalities, betas from ToM engagement had a larger spatial pattern correlation with meta-analytic maps (all *t*(89)s > 5.6, all *p*s < .001; Figure 4c). This analysis could not be performed on the ToM group maps (Dufour et al., 2013) because *t* values were not available for all voxels. This indicates that the neural activity associated with ToM engagement is more similar to existing results about ToM. Together, there is strong evidence that ToM engagement has larger statistical power at localizing ToM-related brain regions than ToM demands.

### Comparison between social interaction perception and theory of mind

So far, we have shown that the group-level summaries (medians) of online participants’ ratings can serve as a reliable and sensitive measure to find the neural correlates of social interaction perception and ToM in the narratives. The medians of the two ratings were correlated to a low to moderate degree (*r* = .29 for Text modality, *r* = .37 for Audio modality, *r* = .32 for all time points; Figure 2c). This supports the view that people tend to use ToM at the same time as perceiving social interactions during natural experience and there may not be a complete distinction between the two processes on the behavioral level. Based upon that, we sought to investigate the common and distinct neural correlates of social interaction perception and ToM.

#### Subheader 1. Contributions of both social interactions and ToM to neural activity in overlapping regions

We fitted a GLM with both the medians of self-reported social interactions and the medians of self-reported ToM, with separate regressors for Text and Audio modality. The correlations between social interaction and ToM ratings stayed relatively constant after being convolved with the canonical hemodynamic response function (*r* = .29 for Text modality; *r* = .38 for Audio modality). Those correlations did not raise concerns for multicollinearity problems (variance inflation factors < 1.3 for all regressors in the model).

Between the Text and Audio modalities, the effect maps of social interaction perception when controlling for ToM were not as similar as when being modeled alone (*r* = .50 for unthresholded *t* maps, Figure 5a left); similar for the effect maps of ToM (*r* = .51, Figure 5b left). Still, we took the conjunction across modalities (Figure 5a and 5b right) because the common neural correlates across modalities were more invariant to low-level stimulus features and thus more specific to the mental processes to be compared. The brain regions responding to social interactions under both modalities include bilateral PC, STS, IFG, dmPFC, and left TPJ; those responding to ToM include bilateral TPJ, PC, IFG, dmPFC, and left STS, as well as bilateral premotor cortex (PMC), intraparietal sulcus (IPS), and bilateral lateral occipitotemporal cortex (LOTC) which encompasses the middle temporal and medial superior temporal area (MT+). Besides whole-brain analysis, we tested the effects of social interactions and ToM in some selected cortical regions (Glasser et al., 2016; Table 2), namely left TPJ, bilateral dmPFC, right PMC, bilateral MT+ (Figure 5c), bilateral PC, left STS, right TPJ, and bilateral supplementary motor area (SMA) (Figure S5). In some selected regions like left TPJ, both social interactions and ToM had significant effects under both modalities, while in some other regions like right PMC, only ToM had significant effects. These results imply that social interactions and ToM have largely overlapping neural correlates, and both contribute independently to driving the neural activity in those regions. At the same time, ToM appears to recruit brain regions that social interactions do not.

**Figure 5.**
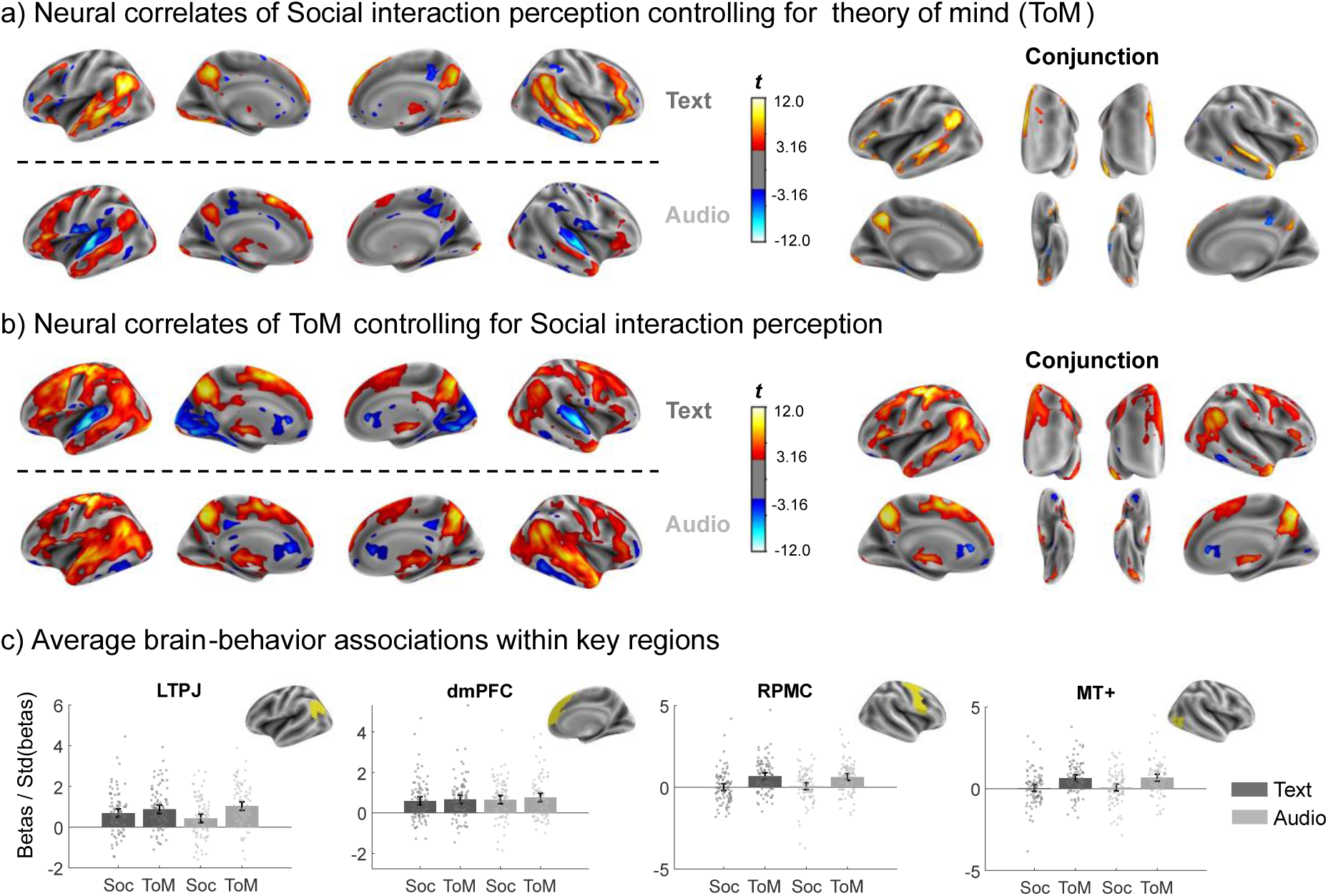
Neural activity associated with social interaction perception and theory of mind (ToM). a) and b) Group-level effect maps of medians of social interaction and ToM ratings after controlling for each other, thresholded at FDR *q* < .01. The method for projection from volumetric space to surface space was the same as Figures 3a and 4a. Surfaces on the left show neural correlates under Text and Audio modality separately, while surfaces on the right show the conjunction across modalities. The color bar of *t* values applies to all surfaces. c) Bar plots of the brain-behavior associations in four regions of interest. Bar heights indicate the means of first-level betas divided by their standard deviation, error bars indicate 95% confidence intervals, and shaded scatters indicate estimates for each participant. Abbreviations: std – standard deviation, LTPJ – left temporoparietal junction, dmPFC – dorsomedial prefrontal cortex, RPMC – right premotor cortex, MT+ – middle temporal and medial superior temporal area, Soc – social interactions.

#### Subheader 2. Cross-modal neural correlates of social interaction perception as a subset of those of ToM

We quantitatively investigated which voxels and regions showed significant correlations with both social interactions and ToM, or only one of them but not the other. To do so, we calculated Bayes Factors for each voxel and region of interest to quantify the evidence for the alternative and null hypothesis, or presence and absence of an effect. For one voxel or region, if we had significant evidence for the alternative hypothesis of the effects of both ToM and social interactions, we would classify it as “ oth”; if we had significant evidence for the alternative hypothesis of ToM and the null hypothesis of social interactions, we would classify it as “ToM only”; in the opposite case, we would classify it as “ ocial interactions only”. We chose 5 as a threshold for the alternative hypothesis and 1/5 for the null hypothesis to obtain moderate to strong evidence given the current sample size (Rouder et al., 2009).

As seen in Figure 6a, across two modalities, both social interactions and ToM correlated with neural activity in bilateral PC, dmPFC, STS, IFG, and left TPJ. In Text modality, social interactions showed unique correlations with activity in right STS, left early visual cortex, and bilateral ITC; in Audio modality, right early visual cortex and left PMC. However, social interactions showed rarely any unique correlations that were common across modalities. By contrast, ToM showed cross-modal unique correlations with neural activity in left anterior IPS, right PMC, bilateral MT+, and left primary somatosensory cortex. We calculated the mean betas and Bayes Factors in the same eight selected regions as for the beta maps and found further evidence that there were regions responding to “ oth” and regions responding to “ToM only” consistently across modality, but not for “ ocial interactions only” (Figure 6b and Figure 5). cross two modalities, activity in about 6.46% of all voxels was correlated with both social interactions and ToM, about 2.84% with ToM only, and a negligible proportion (< .01%) with social interactions only (Figure 6c). The results provide strong evidence that the common neural correlates of social interaction perception across modalities are a subset of those of ToM during naturalistic experience. Besides, several brain regions are only associated with ToM but not social interactions.

**Figure 6.**
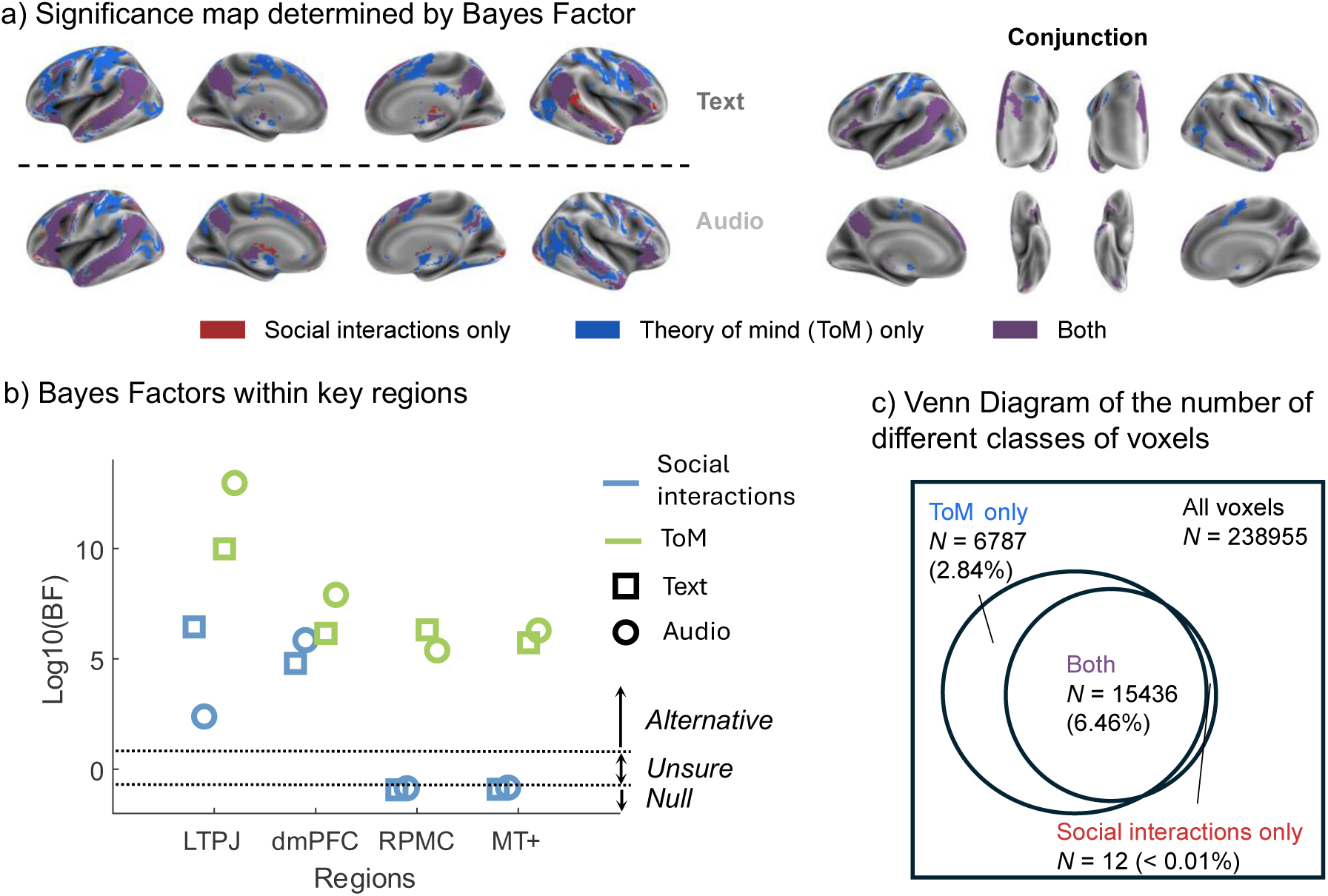
Classifications of voxels/brain regions by Bayes Factors. a) Surface projection of the class of each voxel (Social interactions only, ToM only, or Both). Surfaces on the left show results for Text and Audio modality separately, and surfaces on the right show the conjunction across modalities. The surface space was the same as those in previous figures but the projection method was nearest neighbor here. b) Group-level Bayes Factors for the same four regions shown in Figure 5c (also with the same abbreviations). The thresholds applied to Bayes Factors were shown by horizontal black dashed line, and the corresponding conclusions in each of the three Bayes Factor value ranges were named by italic notes on the right. For example, if one data point has a Bayes Factor larger than 5 (Log10(BF) > ∼0.7), there is sufficient evidence supporting the alternative hypothesis that an effect is significantly above zero. c) A Venn diagram demonstrating the proportions of voxels that belonged to each class in the conjunction map as shown in Panel a. The percentages in parentheses refer to the ratios of the number of voxels in one class over the number of all voxels in the brain.

## Discussion

Using a combination of normative subjective ratings and naturalistic neuroimaging during the experience of written narratives, we demonstrated that in closer-to-real-life experiences, perceiving social interactions and using theory of mind (ToM) co-occurred to a small to moderate extent. They had both common and distinct neural correlates across sensory modalities. In a side analysis, we showed that self-report measurement was a valid and potentially better way to study neural correlates of ToM because it was more sensitive than the annotation method widely adopted by previous studies (e.g., Lee Masson & Isik, 2021).

In the remainder of the Discussion, we will compare our results with previous findings and seek a new interpretation beyond the processing-stage theoretical framework. Specifically, we argue that social interaction perception should be construed as a combination of perceptual and conceptual processes, that both social interaction perception and ToM involve pre-reflective inferential processes, and that ToM additionally involves deliberative inferences. We will unpack these points in more detail and consider the limitations of the current report and potential future work.

### Subheader 1. Social Interaction Perception Involves Both Perceptual and Conceptual Processing

There is rich evidence about the neural correlates of visual social interaction perception and specific visual features embedded in social interactions, such as action directions (Wurm & Caramazza, 2019) and motion information (Landsiedel et al., 2022). Only recently have researchers noted the contribution of multimodal processing to neural activity underlying social interaction perception (Landsiedel & Koldewyn, 2023; Lee Masson et al., 2024b). In the current study, we extended this line of work by showing that verbal descriptions of social interactions engaged similar brain regions to visual and audiovisual presentations. The neural correlates of social interaction perception were also highly similar across the text and audio modalities, providing strong evidence that social interactions in narratives are processed in a modality-general way.

The neural correlates of social interaction perception shared across modalities included some largely visual regions such as LOC, ITC, and early visual cortex (Glasser et al., 2016). This held true even under the audio modality when there were no visual stimuli at all (Figure 3a), which suggests that perceiving social interactions in narratives partly relies on lower-level perceptual processing. Additionally, the neural correlates included many multimodal brain regions such as TPJ, STS, PC, dmPFC, IFG, and medial frontal gyrus, which are key regions involved in mentalizing (Schurz et al., 2021). They also constitute the default mode network (DMN) (Smallwood et al., 2021; Yeo et al., 2011) that has been shown to encode higher-level conceptual information during narrative experiences (e.g., Chen et al., 2017; Simony et al., 2016; Zadbood et al., 2022). This indicates that social interactions in narratives are processed in part as high-level amodal, conceptual information, and they might be an important content that people encode during narratives (Lee Masson et al., 2024b). Given the modality-general findings about its neural correlates, social interaction perception should be more than just an early perceptual process but also involve later conceptual processing.

### Subheader 2. Social Interaction Perception and ToM Both Involve Pre-Reflective Inferences

There has been a popular view that perceiving social interactions naturally invites the engagement of ToM and that the presence of social interactions improves people’s ability to mentalize (e.g., Cross et al., 2016; Dziobek et al., 2006; Walter et al., 2009). However, few studies have directly tested the covariance between social interactions and ToM. By adopting the self-report method, we showed that people’s tendency to use ToM positively correlated with their judgments of social interactions to a small to moderate extent. Additionally, we found that almost all brain regions responding to social interactions across modalities also responded to ToM. Because we included both social interactions and ToM in the same model and they were behaviorally separable, it is unlikely that the neural correlates of social interaction perception could be fully explained by ToM. Instead, there should be computations shared between social interaction perception and ToM that are implemented in their overlapping neural correlates.

The common neural correlates of social interaction perception and ToM included TPJ and dmPFC, which are implicated in pre-reflective, coherent, effortless experience (Lieberman, 2022). They also included other regions in the DMN whose homologous regions in rhesus monkeys have selective responses to social interactions (Sliwa & Freiwald, 2017). Given this cognitive automaticity and evolutionary primacy, the shared neural correlates may function to serve automatic, pre-reflective social inferences. The goal of these inferences is to simulate the social world around us (Mitchell, 2009) and prepare ourselves to act in it. The inferences could be engaged whenever we need to make sense of an agent outside the self, here, and now, whether for understanding social interactions and others’ minds, disambiguating animations (Nguyen et al., 2019), or recollecting autobiographical memories (Rabin et al., 2010). This view argues against the processing-stage framework, indicating that ToM may involve more automatic processing than previously thought and that social interaction perception and ToM involve computations on the same processing level.

### Subheader 3. ToM Involves Deliberative Inferences Limited by Cognitive Resources

Beyond the canonical ToM regions shared with social interaction perception, ToM engagement ratings in the current study were associated with neural activity in other regions, including the right anterior part of IPS, left PMC, and bilateral LOTC or MT+, which were less commonly reported in existing studies (Schurz et al., 2021). We interpret these results as implying that ToM involves deliberative operation and manipulation over mental-state-related information. Specifically, the anterior part of IPS and PMC are important in understanding the goals and intentions of actions, a key feature of the “mirror” system (Zaki & Ochsner, 2009). The mirror system can transmit information to mPFC and lateral temporal cortex to serve higher-order mentalizing, especially when people deliberate about mental states from simpler perceptions (e.g., de Waal & Preston, 2017; Van Overwalle & Baetens, 2009). Similarly, LOTC is sensitive to action verbs (Taylor et al., 2017) and possible actions (affordances) towards a visually presented tool (Wu et al., 2020). Thus, the association between neural activity in PMC, anterior IPS, and LOTC and ToM engagement may reflect voluntary efforts for mentalizing from the action cues in narratives.

Importantly, the controlled mental processing unique to ToM should be different from pre-reflective inferences and gated by limited cognitive resources. Two observations support this view: First, the IPS and PMC are also important for executive functions, such as multitasking, which entail deliberation and reflection (Worringer et al., 2019). Second, ToM might have “recycled” e isting neural infrastructure for social interaction understanding (especially the DMN) during evolution to support its automatic, core functioning (Sliwa & Freiwald, 2017). Considering that humans are capable of abstract inferences based on formal propositions but nonhuman primates are not (Deen et al., 2023), the unique ToM neural correlates outside the DMN should be supporting more cognitively demanding inferences. Thus, our results imply that ToM includes both pre-reflective and controlled inferences, with the former engaged by perceiving social interactions but not the latter.

### Subheader 4. Limitations and Future Directions

Some limitations of the current study are worth mentioning. First, most of our results were based on group-level statistical inferences. Although this provided insights into the “on average” effects, there must be important individual differences omitted, especially considering the idiosyncratic nature of ToM. More work is needed in developing new ways to measure ToM engagement and relate it to neural activity on the individual level. Second, given the limited spatial resolution of fMRI, we may have missed some brain regions that only respond to social interactions across modalities. It will be helpful to investigate the dose-response profiles of the regions related to social interactions and ToM to get a finer picture of how they process the two types of information. Third, our results and interpretations do not consider action understanding as a separate psychological construct on its own, although it has shared and unique neural correlates from social interaction perception and ToM (Arioli & Canessa, 2019). Some of our results are thus likely to be attributed to action understanding. Future work should model social interactions and ToM together with actions in the stimuli to get a finer picture of their neural correlates.

## Conclusions

The current study shows that the modality-invariant processing of social interactions relies on a subset of the brain regions responding to ToM in narratives, while ToM has its unique neural correlates. This implies that both perceiving social interactions and ToM involve pre-reflective inferences, with social interaction perception having its own perceptual components and ToM having its own deliberative inference components. Future research is needed to test the reliability and generalizability of these findings and to refine the theoretical accounts for them.

## Methods

### Behavioral ratings of social interactions and theory of mind

#### Subheader 1. Participants

A total of 268 participants took part in our online experiment from Prolific.com. Of these, 32 participants failed more than half of the attention check questions, two participants provided the same ratings throughout the experiment, and three participants did not have their data saved. The remaining 231 participants with valid data (104 females, 127 males) are all healthy adults (19-45 years old, *M* = 33.65, *SD* = 6.51) whose native language is English and who have no literacy difficulties. All participants have an approval rate (proportion of valid data from studies they have taken part in) of over 95% on Prolific.

#### Subheader 2. Stimuli

The main stimuli are eight short narratives (stories) in the third-person perspective. Each narrative includes a single protagonist who moves along a single storyline composed of nine different situations, which are sampled from Polti’s 6 dramatic situations (Polti, 1917) and characterized by interpersonal relationships and actions. During the e periment, each situation or “part” was presented in one e perimental trial. The first four narratives were presented as audios, where an AI-generated adult male voice read out the narratives. The second four were presented as white texts on a black full-screen background at a constant speed of three words per second; the number of words on each screen was 10 or the number of remaining words in a part, whichever was smaller (Figure 1a). Table 1 below provides a more detailed description of the duration and length of each narrative, the presenting modality, and the duration in the unit of volumes (TRs) in the neuroimaging study.

**Table 1.** Properties of the narrative stimuli.

| Run # | Narrative # | Modality | Duration (mm:ss) | Number of TRs | Number of words |
| --- | --- | --- | --- | --- | --- |
| 1 | 7 | audio | 2:01 | 263 | 429 |
| 1 | 8 | audio | 1:36 | 209 | 314 |
| 2 | 5 | audio | 2:30 | 326 | 511 |
| 2 | 6 | audio | 2:04 | 269 | 447 |
| 3 | 3 | text | 3:11 | 415 | 573 |
| 3 | 4 | text | 3:29 | 454 | 626 |
| 4 | 1 | text | 2:43 | 355 | 490 |
| 4 | 2 | text | 2:40 | 348 | 480 |

*Note.* The table is sorted by run number followed by narrative number. The sequence of the two narratives within a run was counterbalanced across participants.

Importantly, the variety of situations and transitions between them make each narrative contain rich descriptions of both the activity of a single character and the joint activity of multiple characters, allowing for a contrast over social interactions. Similarly, some parts of the narratives contain dense descriptions of characters’ mental states (e.g., “she was furious that she did not know what was going on”) while others do not (e.g., “ my and Linda continued to be best friends”), making it possible to induce different levels of ToM engagement at different times during the narratives.

#### Subheader 3. Experimental Procedures

Potential participants open the study link from Prolific.com and will be directed to the webpage-based experiment hosted on Pavlovia.org. There they learn more about the experiment and provide their consent of participation before starting the experiment.

The experiment consists of four runs. Between two runs, participants could take a short break of no more than one minute. In each run, two narratives are played as texts or audios (Table 1). Because each narrative is presented in nine experimental trials, there are 18 trials in each run. At the start of the first and third runs, there is an additional practice trial for participants to get familiar with the experimental interface and ratings. One similar but irrelevant narrative part is used in the practice trials.

Each trial starts with a fixation period with pseudo-randomized durations between 2 and 8 seconds. Then a narrative part is presented, while participants need to give their ratings moment by moment by moving their mouse on top of a continuous visual analogue scale with two anchors at the ends and one anchor at the midpoint. The rating question is randomly chosen for each participant. About half of the participants have answered the social interaction question: “*Are there social interactions? (How much do you think there are social interactions in the narrative at this moment?)*”, with three anchors being “*Not at all*”, “*Partially*”, and “*Definitely yes*”. The other half of the participants have answered the theory of mind question: “*Thinking about mental states? (How much are you thinking about the mental states of characters in the narrative at this moment?)*”, with three anchors being “*Not at all*”, “*Half the time*”, and “*All the time*”. Questions in the bracket are only presented in practice trials. t the end of each trial, participants are asked to give an “intermittent” rating about the previous trial: “*On average, how much [were you thinking about the mental states of characters]/[do you think there were social interactions] during the last part of the narrative?*” The anchors are the same as the previous ratings (Figure 1). The intermittent ratings have not been analyzed or reported in this article.

The sequence of the two narratives within each run is counterbalanced across participants. To check whether participants were paying attention, in one random trial per run, a surprise question would follow the intermittent rating and ask participants to click on a specific location of the scale.

#### Subheader 4. Data collection

All ratings were recorded as a float number between 0 and 100, linearly mapping onto the positions of the visual analogue scale. The moment-by-moment ratings were sampled every ms or every cycle of the participants’ monitor refreshing, whichever was longer. ecause we anticipated a delay in participants’ ratings as compared to the narrative flow (e.g., a “yes” rating to social interactions might happen hundreds of milliseconds after social interactions appeared), we continued collecting ratings for another 1.5 seconds after each narrative part ended.

As stated in the Results section, researchers annotated ToM demands in the narratives offline. Because researchers did not experience the narratives continuously as participants did, only “yes no” (1 0) labels were used for annotations, instead of continuous values. When there was a conflict of annotations, that is when two raters provided the same label while the other two opposed, the consensus label was generated after further discussion. 24 out of 172 (14%) sentence parts had a conflict of annotations.

#### Subheader 5. Quantification and statistical analysis

To ensure data quality, we excluded ratings of a participant from further analysis if they failed more than two out of the four attention check questions (*n* = 32) or kept providing the same rating throughout the experiment (*n* = 2). During the first few seconds of each trial, participants might not have started engaging in the narrative and providing meaningful ratings. Thus, we set as missing values all the moment-by-moment ratings sampled before the first move of the mouse in each trial. Next, to aggregate and compare moment-by-moment ratings across participants, we down-sampled them to 230ms per sample (chosen as half of the resolution of the BOLD data in the neuroimaging study) by linear interpolation. In generating a group-level summary statistic that was later applied to neuroimaging data, we excluded data from participants whose social interaction ratings had a correlation with the medians of all other participants’ ratings lower than 0. (*n* = 7), as well as those whose ToM ratings had a correlation lower than 0.2 (*n* = 14).

To estimate the extent to which online participants had similar ratings for social interactions and ToM, two metrics were calculated. First, we randomly divided participants into two equal-size groups and correlated the mean ratings of the two groups. This split-half correlation value was corrected by the Spearman-Brown formula (Brown, 1910; Spearman, 1910). We repeated the sampling and calculation 2000 times randomly to estimate the mean and confidence interval of the correlation coefficients. Second, we calculated the pairwise correlation between participants’ ratings and found the range, mean, and median of the correlations. The latter metric is a more conservative estimate of how similar participants were in their ratings.

We also investigated the e tent to which researchers’ annotations of ToM demands agreed with online participants’ ratings of ToM engagement. To do that, we resampled the annotations to 230ms per sample by linear interpolation. We then extracted the medians of online participants’ ratings as a group-level summary and calculated its correlation with researchers’ annotations. The statistical significance of the correlation was estimated by 10000 bootstrap samples with phase randomization (Lancaster et al., 2018).

### Neuroimaging study on social interaction perception and theory of mind

#### Subheader 1. Participants

Ninety-seven participants took part in an fMRI experiment. Data from one participant was missing due to technical issues, and data from another six participants were excluded from further analysis because of failures in fieldmap collections, which are essential for the susceptibility distortion correction step in fMRI preprocessing. The remaining 90 participants with valid data (55 females, 34 males, 1 other) are healthy adults (18-45 years old, *M* = 24.72, *SD* = 5.57) without any current physical or mental disorders. All participants are native English speakers or have comparable fluency in English, and have normal or corrected-to-normal vision.

#### Subheader 2. Stimuli and Experimental Procedures

The neuroimaging study is part of a larger project about the neural correlates of various cognitive and affective processes (Jung et al., 2024). The project was conducted at the Dartmouth Brain Imaging Center and required participants to pay four separate visits on different days. ata analy ed here came from the “narratives” task, which was performed on the second visit of every participant. At the beginning of their second visit, participants provided their written consent to continuing participation in the study. After that, they practiced every task they would perform that day on a laptop under e perimenters’ instructions. ll instructions were previously scripted and read out loud to each participant. When participants had no further questions about the tasks, they would lie into the scanner and begin the formal tasks.

The run structure and narratives played in each run were the same as in the behavioral study (Table 1). The main difference between the two studies was in the trial structure: fMRI participants did not provide any ratings when the narratives were presented; after one narrative part finished, they were asked to provide two ratings about their feelings and expectations. The ratings were not analyzed or reported in the current article (Figure S1).

#### Subheader 3. Data collection

Structural and functional MRI data were acquired on a 3T Siemens MAGNETOM Prisma MRI scanner with 32-channel parallel imaging. Structural images were acquired using high-resolution T1 spoiled gradient recall images. Functional images were acquired with a multiband EPI sequence (repetition time (TR) = 460ms, echo time (TE) = 27.2ms, field of view = 220mm, multiband acceleration factor = 8, flip angle = 44°, 64×64 matrix, 2.7×2.7×2.7mm voxels, 56 interleaved ascending slices, phase encoding posterior to anterior). For more details about the scan, see Jung et al. (2024). Stimuli were programmed and presented by Psychtoolbox (MATLAB, MathWorks) on a Linux laptop.

#### Subheader 4. Quantification and statistical analysis

##### MRI data preprocessing

Preprocessing was performed using fMRIPrep 21.0.2 (Esteban et al., 2019) based on Nipype 1.6.1 (Gorgolewski et al., 2011). To correct for magnetic field inhomogeneity, a B0-nonuniformity map (fieldmap) was estimated based on two echo-planar imaging (EPI) references with `topup` (Andersson et al., 2003, FSL 6.0.5.1:57b01774).

For each participant, one T1-weighted (T1w) image was corrected for intensity non-uniformity with `N4BiasFieldCorrection` (Tustison et al., 2010), distributed with ANTs 2.3.3 (Avants et al., 2008, RRID:SCR_004757), and used as T1w-reference throughout the workflow. The T1w-reference was then skull-stripped with a `Nipype` implementation of the `antsBrainExtraction.sh` workflow using OASIS30ANTs as the target template. Brain tissue segmentation of cerebrospinal fluid (CSF), white-matter (WM) and gray-matter (GM) was performed on the brain-extracted T1w using `fast` (Zhang et al., 2001, FSL 6.0.5.1:57b01774, RRID:SCR_002823). Brain surfaces were reconstructed by `recon-all` (Dale et al., 1999, FreeSurfer 6.0.1,RRID:SCR_001847). The brain mask estimated previously was refined with a custom variation of the method to reconcile ANTs-derived and FreeSurfer-derived segmentations of the cortical gray-matter of Mindboggle (Klein et al., 2017, RRID:SCR_002438). Volume-based spatial normalization was performed through nonlinear registration with `antsRegistration` (ANTs 2.3.3), using brain-extracted versions of both T1w reference and the T1w template. Two templates were selected for spatial normalization: *ICBM 152 Nonlinear Asymmetrical template version 2009c* (MNI152NLin2009cAsym, RRID:SCR_008796; TemplateFlow ID: MNI152NLin2009cAsym) and *FSL’ M I ICBM 152 o -linear 6th Generation Asymmetric Average Brain Stereotaxic Registration Model* (RRID:SCR_002823; TemplateFlow ID: MNI152NLin6Asym).

For each of the four BOLD runs per subject, the following preprocessing was performed. First, a reference volume and its skull-stripped version were generated by aligning and averaging 1 single-band reference (SBRefs) using a custom method of fMRIPrep. Head-motion parameters with respect to the BOLD reference (transformation matrices, and six corresponding rotation and translation parameters) are estimated before any spatiotemporal filtering using `mcflirt` (Jenkinson et al., 2002, FSL 6.0.5.1:57b01774). The estimated fieldmap was then aligned with rigid registration to the target EPI reference run. The field coefficients were mapped on to the reference EPI using the transform. The BOLD reference was then co-registered to the T1w reference using `bbregister` (FreeSurfer) which implements boundary-based registration (Greve & Fischl, 2009), generating a preprocessed BOLD run in MNI152NLin2009cAsym space. Co-registration was configured with nine degrees of freedom to account for distortions remaining in the BOLD reference. Several confounding time series were calculated based on the preprocessed BOLD: framewise displacement (FD), DVARS and three region-wise global signals. FD and DVARS were calculated for each functional run both using their implementations in `Nipype` following the definitions by Power and colleagues (Power et al., 2014). The three global signals were extracted within the CSF, the WM, and the whole-brain masks. The six head-motion estimates were expanded with the inclusion of temporal derivatives and quadratic terms for each (Lund et al., 2006). Gridded (volumetric) resampling was performed using `antsApplyTransforms` (ANTs), configured with Lanczos interpolation to minimize the smoothing effects of other kernels (Lanczos, 1964). Lastly, the functional data were spatially smoothed with a 6 mm full-width-at-half-maximum (FWHM) Gaussian kernel.

##### Voxel-wise general linear model (GLM)

To find the neural correlates of variables of interest (social interactions and ToM), we applied a hierarchical random effect model approach using two-stage summary statistics. Specifically, in each voxel, we fit a GLM for each participant using the ordinary least squares method. We then aggregated the regression coefficients in each voxel across participants and performed one-sample t-tests and robust regression (Holland & Welsch, 1977) on single coefficients or planned contrasts over several coefficients. Results from t-tests and robust regression were almost identical and we only reported those from t-tests. To find labels for the significant brain regions on the cortex, we overlaid the thresholded *t* maps on a multi-modal parcellation of the human cerebral cortex (Glasser et al., 2016) and calculated the percentage of significant voxels within each parcel. Parcels with more than 25% of significant voxels were treated as significant, and adjacent parcels were grouped into larger regions with more commonly used names (such as TPJ instead of PGi and PGs).

Variables of interest came from two different sources: online participants’ moment-by-moment ratings and researchers’ offline annotations. The medians of online participants’ ratings were linearly pro ected from the range [0, 100] to [-1, 1] before being entered as regressors in the GLM. To ensure the medians were valid and meaningful, for each trial, we excluded data from participants who had less than 40% of non-missing moment-by-moment ratings. We also excluded the time points when less than 50% of the participants had non-missing rating values by adding indicator functions to the GLM whose values were 1 at the e cluded time points and 0 at all other time points. Researchers’ annotations of ToM demands were coded as two separate binary regressors with values 1 or 0. One regressor modeled the sentence parts where there were ToM demands and the other modeled where there were no ToM demands; for a sentence part with ToM demands, the first regressor had a value of 1 and the second a value of 0. Thus, a contrast between the estimates of the two regressors revealed the effects of ToM demands.

To control for the effect of answering questions on neural activity, we added a binary regressor whose values were 1 when participants were giving ratings at the end of each trial and 0 otherwise. This rating regressor, together with all variables of interest, was convolved with the canonical hemodynamic response function (Glover, 1999). This made the fixation periods the only implicit baseline of the BOLD signals.

Some further data denoising was performed in the GLM. First, we high-pass-filtered the functional data at 1/128 Hz by adding a series of discrete cosine functions to each model. Second, we added 24 head-motion-related parameters to account for the effects of head motions, including six head motion parameters (x, y, and z translations, and pitch, yaw, and roll), their temporal derivatives, their squares, and the squares of the derivatives. Third, we added the mean signal in the cerebral-spinal fluids as an approximation of physiological artifacts. Lastly, we identified outlier brain volumes by searching for the volumes whose average signals or framewise displacements were outside three standard deviations from the mean of each run. We excluded those volumes by entering each of them as an indicator function whose values were 1 at the time point of the outlier and 0 at all other time points.

Because the functional signals at the start of each run may be unstable and deviate from the rest of the run, we discarded the first 6 volumes in each run when there were no stimuli on the screen. The rest of each run’s functional data were rescaled to have a grand mean of 100 (across all volumes and voxels) to account for differences in signal baselines across runs and participants (Chen et al., 2017). Functional data were concatenated across all four runs, as were the design matrices. Variables of interest, as well as the rating regressor, were shared across runs, such that only one beta value was estimated for each variable of interest in each voxel. All other variables, including the intercept, were modeled separately for each run.

##### Mask-based analysis

To understand whether our fMRI results were congruent with existing findings, we performed analysis within several brain masks defined by results from previous studies. The first way to find brain masks was to use term-based association maps from Neurosynth.org (Yarkoni et al., 2011), whose z-scores from a two-way ANOVA quantify the possibility of each voxel being positively activated in a paper that includes a specific term. The original maps were thresholded at FDR *q* < .01, and we further thresholded them by *z* >= 3 to reduce false positive voxels and make the masks more specific. The terms used in the current study include: “social interaction”, “social cognition”, “psts”, “tom”, and “mentalizing”.

The second way to find brain masks was to use the ToM group maps from a false belief task (Dufour et al., 2013). The masks include bilateral TPJ (combining the original “LTPJ” and “RTPJ” maps), RSTS(the original “RSTS” map), bilateral PC(the original “PC” map), and bilateral dmPF (combining the original “DMPFC” and “MMPFC” maps). We did not use the “DMPF” map because no GLM results in the current study showed significant effects in the ventral medial prefrontal cortex.

The third way was to define region masks based on a multimodal parcellation of the cerebral cortex (Glasser et al., 2016). Each mask was a union of several parcels based on their anatomical positions and functional topography. For a full list of masks used in the current study and how we defined them, see Table 2. Within each mask defined in any of the three ways, we extracted individual-level regression coefficients from the GLM analysis and compared the group mean to zero to investigate whether a variable of interest had consistent neural activations across participants in that mask.

**Table 2.** Masks defined on parcellations from Glasser et al. (2016)

| Mask | Parcels included | Hemisphere |
| --- | --- | --- |
| LTPJ | PGi, PGs | Left |
| dmPFC | d32, 8BM, 9m | Bilateral |
| RPMC | FEF, PEF, 6d, 6v, 6r, 6a | Right |
| MT+ | V4t, FST | Bilateral |
| PC | 7m, v23ab, d23ab, 31pv, 31pd | Bilateral |
| LSTS | STSda, STSdp, STSva, STSvp, TPOJ1, TPOJ2 | Left |
| RTPJ | PGi, PGs | Right |
| SMA | SCEF, 6ma, 6mp | Bilateral |

*Note.* A “Bilateral” entry in the Hemisphere column denotes that the parcels on both hemispheres were concatenated to form a single brain mask for analysis. “Left” and “Right” denote that parcels on only one hemisphere were include.

##### Bayes Factor analysis

To identify voxels associated with both social interactions and ToM, and those associated with one but not the other, we took the Bayes Factor (BF) approach to quantify evidence for both alternative and null hypotheses. Specifically, the BF value of a given voxel is proportional to the ratio of the likelihoods under the alternative and null hypotheses (e.g., the neural activity in that voxel is correlated with social interactions in narratives or not). We estimated BFs from *t* values of one-sample tests, assuming a Jeffreys-Zellner-Siow (JZS) prior, following the formula provided by Rouder et al. (2009):

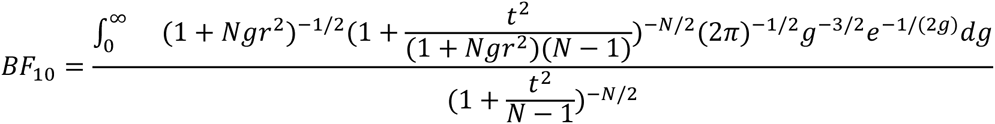

In the formula, *t* represents the *t*-statistic, *N* the sample size, and *r* the scale factor. We set the scale factor to 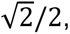 assuming a moderate effect a priori (Rouder et al., 2009), to increase the support for alternative hypotheses given the frequently smaller effect size in fMRI univariate analysis (see also Bo et al., 2024).

BFs calculated in this manner are asymmetric for alternative and null hypotheses, because evidence for null hypotheses is bound by *t* = 0. Given the current sample size (*N* = 90), the BF cannot be smaller than .12 but can be infinitely large. Thus, we chose a heuristic threshold of 5 for the alternative hypothesis and 1/5 for the null hypothesis to attain moderate to strong evidence in favor of both hypotheses.

After calculating the BF for a voxel under one modality, we classified the voxel into one or none of the three categories (Figure 6). Because the BF threshold for alternative hypotheses corresponds to a *t* value of 2.84, which is lower than the critical *t* value (3.16) for FDR *q* < .01, we further bound the alternative results by the significance in the FDR corrected analysis. Besides, we only considered positive effects (positive *t* statistics) in this analysis. The specific criteria are as follows:

- “Both” – BF > 5 and *t* > 3.16 for both Social interactions and ToM.
- “Social interactions only” – BF > 5 and *t* > 3.16 for Social interactions, and BF < 1/5 for ToM.
- “ToM only” – BF > 5 and *t* > 3.16 for ToM, and BF < 1/5 for Social interactions.

To create the conjunction classification map across modalities (Figure 6a), we assigned a label to one voxel in the conjunction map if and only if it had the same label under the two modalities. This provides a more conservative and specific estimate of the modality-general neural system responding to social interactions only, ToM only, or both.

