## Supplementary Materials for "COMMON AND DISTINCT NEURAL CORRELATES OF SOCIAL INTERACTION PERCEPTION AND THEORY OF MIND"

### Supplementary Methods

#### Validation of social interaction perception results

To test the robustness of the neural correlates of social interaction perception, four researchers, including the first author, independently annotated the narratives in terms of social interactions. We annotated one sentence part as having social interactions if and only if it involves more than one character and they have communications or physical interactions. We generated consensus labels in the same way as for ToM demands. In GLM analysis, we again created two binary regressors from the annotations, one for social interactions, the other for no social interactions. A contrast between the estimates of the two regression coefficients revealed the effects of social interactions.

#### Support vector machine (SVM) classification

Modeling the social interaction annotations by two binary regressors allowed us to compare the whole-brain neural correlates of social interactions and no social interactions and test the generalizability of the pattern difference across modalities. Specifically, we performed an SVM analysis where the inputs or features were each participant's whole-brain regression coefficient maps (beta maps) and the outcomes or targets were labels of "social interactions" and "no social interactions". Two linear SVM classifiers were trained for the two modalities separately, with box constraint parameter $C = 1$. Performance of the classifier within each modality was evaluated through five-fold leave-whole-participant-out cross validation (i.e., if the social interaction map of one participant was in the left-out fold, their no social interaction map was also in the left-out fold). Performance across modalities was evaluated by applying the model trained on the entire sample in one modality to the other modality. All classifications took the form of two-alternative forced choices (2AFC), where each time a classifier was presented with two beta maps from one participant and judged which one was the social interaction map. We evaluated classification performance by two metrics: 1) classification accuracy, which is the number of hits/true positives and correct rejections/true negatives over the number of all classifications; 2) *Cohen's d*, which is calculated as the ratio of the mean of the differences of distance from the hyperplane between two beta maps over their standard deviations, and which can serve as an unbiased estimate for effect sizes.

#### Trial progress

As can be seen from Figure S1, for both the fMRI experiment and the online experiment, each experimental trial starts with a fixation period whose duration was selected from a uniform distribution between 2 and 8 seconds and rounded to the nearest integer. The fixation durations were predetermined before the experiment for each trial to ensure the total duration of the fMRI experiment stayed constant. The narrative presentation period was the key time window where all regressors about social interactions and ToM were built. The online experiment had a longer narrative presentation period because we added 1.5 seconds of blank stimuli to the end of each trial, in order for participants to be able to finish their moment-by-moment ratings. The rating period was used to collect ratings about current feelings and expectations about the contents of the narratives in the fMRI experiment, and the average ratings of social interactions or ToM in the online experiment. Ratings in the rating period were not analyzed or reported in the current article.

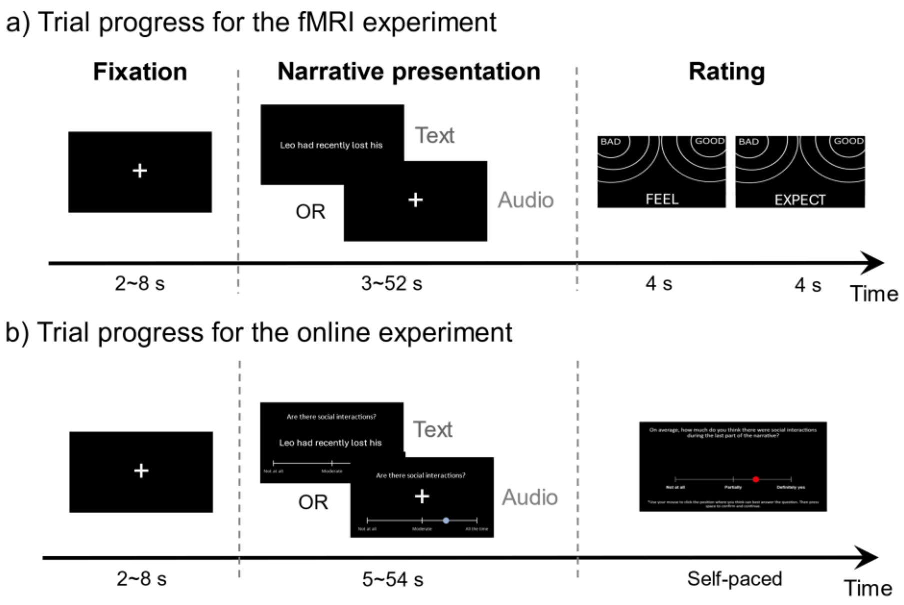


**Figure S1.** Trial progresses for the two experiments. Gray dashed lines indicated boundaries between periods. The period names, Fixation, Narrative presentation, and Rating, applied to both panels.

### Supplementary Results

#### Neural correlates of social interactions by researchers' annotations

The four researchers largely agreed with each other on their annotations about social interactions (pairwise correlations ranged from .66 to .78 (*M* = .71); Fleiss' kappa = .71). In fMRI analysis, we again found very similar effect maps across modalities (*r* = .659; Figure S2a). Across two modalities, the common regions being activated were largely the same as the analysis using online participants' ratings, including TPJ, STS, dmPFC, and so on. Indeed, the unthresholded conjunction maps estimated from the two methods were highly similar (*r* = .870). Those results provided evidence that the neural correlates we found were invariant across measurements, no matter whether they were from researcher annotations or from online participants' ratings.

In the SVM analysis, we found large pattern differences between the "social interaction" and "no social interaction" beta maps, as suggested by the high within-modality accuracies (> 97%) and effect sizes (> 1.8). More importantly, the classifiers trained in one modality also worked well in the other (accuracies > 90%, *Cohen's d* > 1.3, Figure S2b). It indicated that not only general activation patterns but also the differences between patterns caused by social interactions were similar across modalities. Together, those results provided strong evidence that the same neural circuits responded to social interaction information in the narratives both when they were visually and auditorily presented.

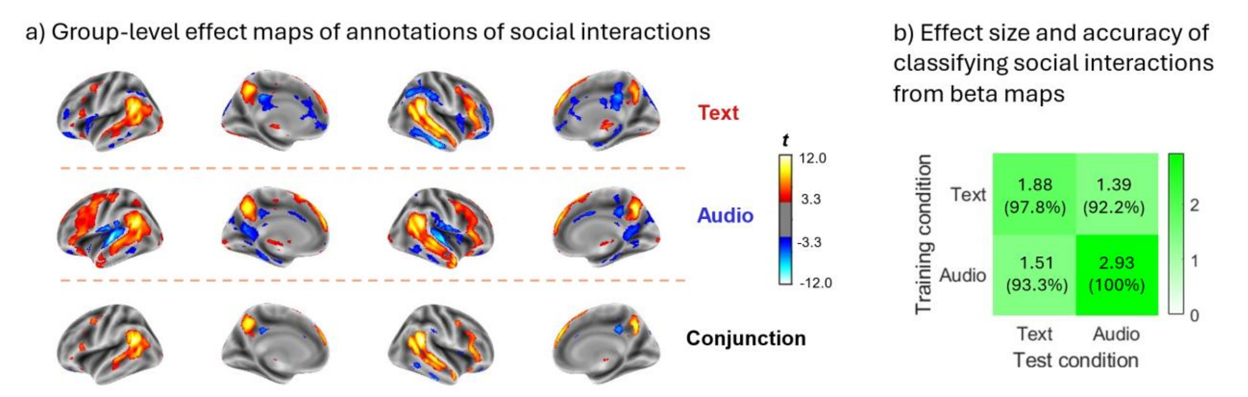


**Figure S2.** Neural correlates of social interactions as researchers' annotations. a) Contrast map of social interaction versus no social interaction. The color bar applies to all surface plots. b) A confusion matrix of a support vector machine (SVM) analysis to predict the labels of whole-brain beta maps (social interactions or no social interactions). Models trained in one modality (Text or Audio) were tested in both modalities. Performance was estimated by classification accuracy *Cohen's d* and reported in the figure (accuracy in brackets).

#### Neural correlates of ToM engagement and ToM demands

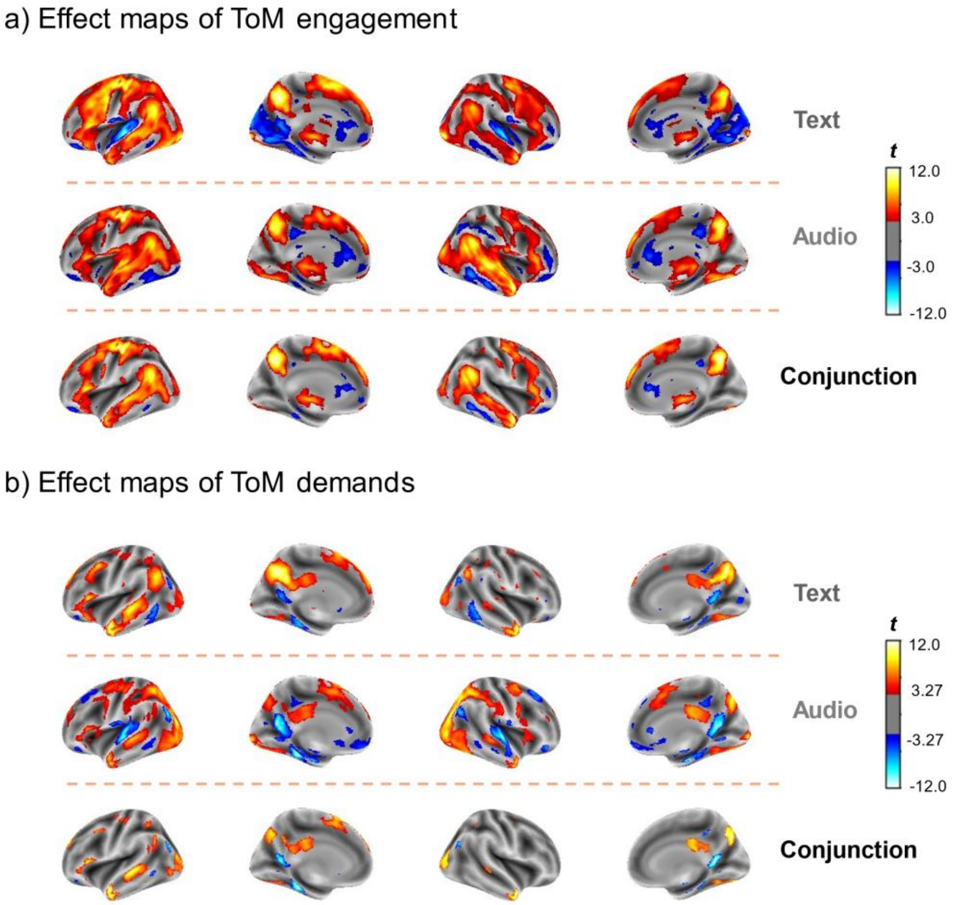


**Figure S3.** Neural correlates of ToM engagment and ToM demands in two modalities separately and their conjunctions. Each set of maps was thresholded at FDR $q < .01$. Across modalities, the neural correlates of ToM were not as similar as social interaction perception but still in the moderate range (on the unthresholded *t* maps, $r = .570$ for ToM engagement, $r = 530$ for ToM demands).

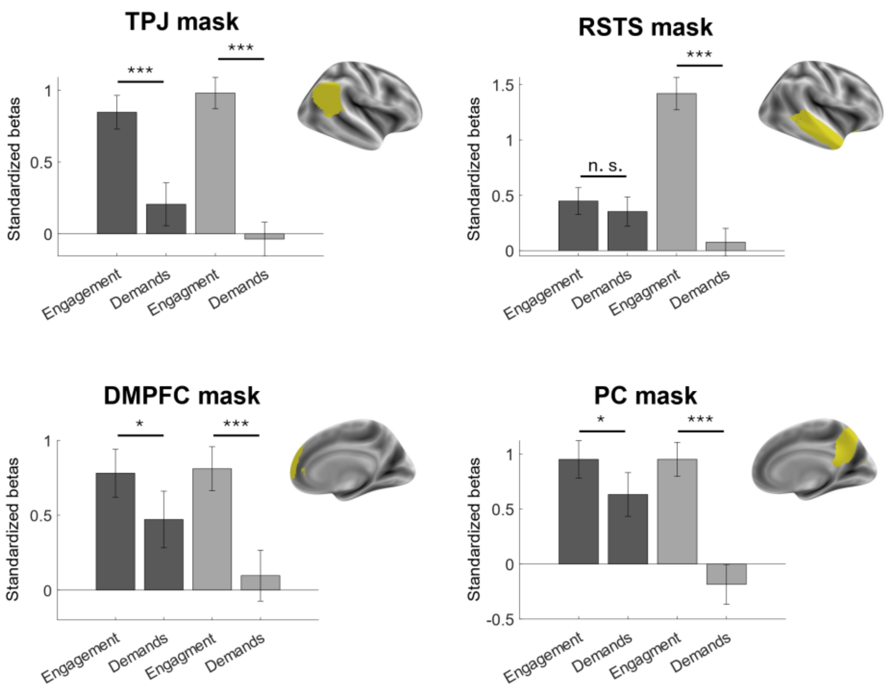


**Figure S4.** Comparison of betas between self-reported ToM and ToM demands. This figure shows comparisons in the brain masks defined by the ToM group map (Dufour et al., 2013; see Methods for details). ***: $p < .001$; *: $p < .05$; n. s.: $p > .05$ (Bonferroni correction applied).

#### Neural correlates of social interactions and ToM

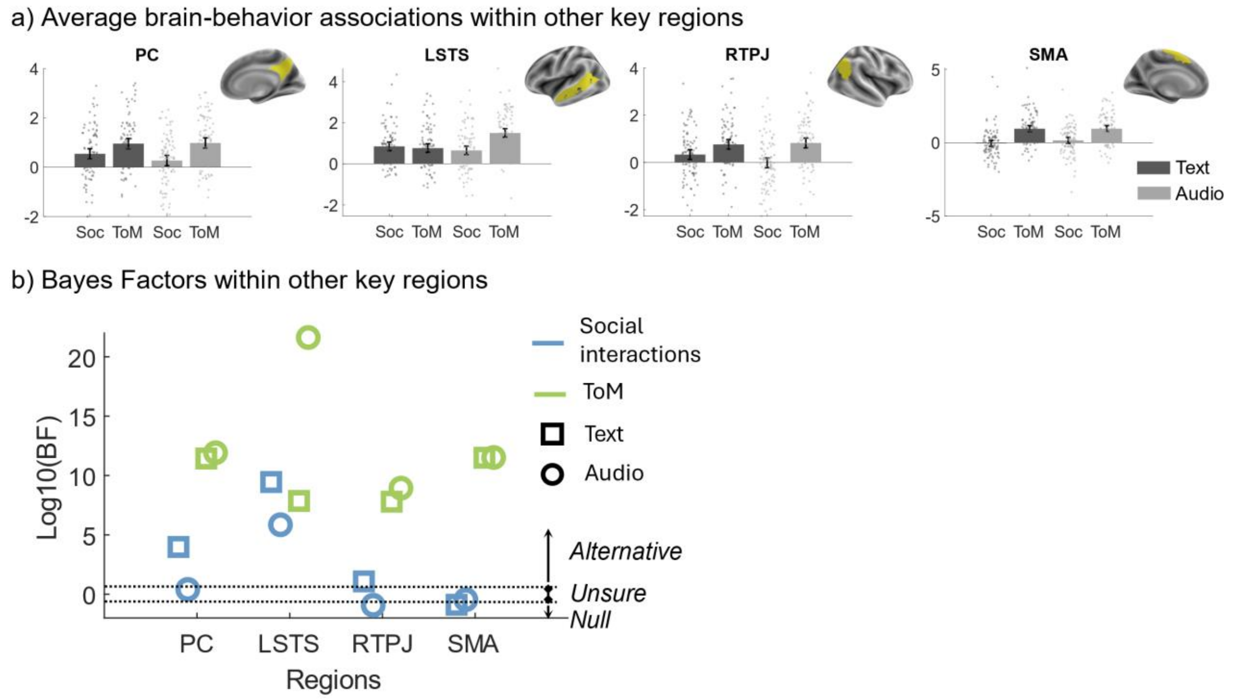


**Figure S5.** Average betas and Bayes Factors in selected brain regions. a) complements Figure 5c and b) complements Figure 6b. All conventions follow those two figures.

#### fMRI effect tables

The tables below summarize the significant clusters/blobs of voxels in the main fMRI analyses reported in the main text. Without extra notes, all tables are organized by one cluster occupying three rows. The first row specifies the volume, peak coordinates (X, Y, and Z separately), and max *t* values. It also specifies which of the 16 cortical "Networks" from the resting-state cortical brain parcellation (Yeo et al., 2011) has largest overlap (quantified by Dice coefficients) with the cluster; an entry of "Sub-cortex" means that the largest proportion of the cluster does not belong to any cortical networks. The third row contains names of the atlas regions which have >= 25% of the voxels with significant effects, with all names from the CANlab 2024 atlas (https://github.com/canlab/Neuroimaging_Pattern_Masks/tree/master/Atlases_and_parcellations/2024_CANLab_atlas). The second row contains "heuristic names" of those region names for easier interpretations, which are commonly used anatomy- or function-based brain region names that overlap with the atlas regions.

**Table S1.** Regions associated with social interactions under both modalities

| Cluster | Volume (mm^3) | X | Y | Z | Max t | Network |
|---|---|---|---|---|---|---|
| 1 | 5416 | 2 | -57 | -45 | 4.67 | Sub-cortex |
| | Cblm cortex R, Cblm cortex L, Cblm vermis | | | | | |
| | *Cblm IX R, Cblm IX L, Cblm Vermis IX* | | | | | |
| 2 | 28896 | -23 | -85 | -17 | 4.17 | Cortex_Visual_Central |
| | visual ventral L, visual early L, visual dorsal L, visual MT+ L, Cblm cortex L | | | | | |
| | *Ctx VMV3 L, Ctx V8 L, Ctx V4 L, Ctx V3CD L, Ctx LO2 L, Ctx V3B L, Cblm CrusII L, Ctx LO1 L, Ctx V3 L* | | | | | |
| 3 | 28496 | 28 | -81 | -19 | 7.03 | Sub-cortex |
| | visual dorsal R, visual ventral R, visual early R, visual MT+ R, Cblm cortex R | | | | | |
| | *Ctx V3CD R, Ctx V8 R, Ctx VMV3 R, Ctx V4 R, Ctx LO2 R, Cblm CrusII R, Ctx LO1 R, Ctx V3 R* | | | | | |
| 4 | 53808 | 56 | -15 | -5 | 7.03 | Cortex_Temporal_Parietal |
| | auditory association cortex R, cingulate vlPFC R, parietal TPOJ R, parietal inferior lobule R, temporal lateral R, cingulate ventral frontal R, cingulate dlPFC R, insula anterior R | | | | | |
| | *Ctx STSvp R, Ctx STSva R, Ctx IFJa R, Ctx 45 R, Ctx TPOJ1 R, Ctx PGi R, Ctx STV R, Ctx TE1a R, Ctx STSdp R, Ctx STSda R, Ctx 44 R, Ctx 47l R, Ctx TPOJ3 R, Ctx IFSp R, Ctx TPOJ2 R, Ctx 8C R, Ctx STGa R, Ctx TGd R, Ctx 47s R, Ctx A5 R, Ctx p9 46v R, Ctx AVI R, Ctx TE1m R* | | | | | |
| 5 | 74712 | -49 | -25 | -1 | 7.03 | Cortex_Default_ModeB |
| | auditory association cortex L, parietal inferior lobule L, parietal TPOJ L, cingulate dlPFC L, cingulate ventral frontal L, somatomotor premotor L, visual MT+ L, cingulate vlPFC L, temporal lateral L, insula anterior L, somatomotor primary L | | | | | |
| | *Ctx STSvp L, Ctx PGi L, Ctx TPOJ2 L, Ctx STSdp L, Ctx STSva L, Ctx 8C L, Ctx TPOJ1 L, Ctx 47s L, Ctx STV L, Ctx 6a L, Ctx MST L, Ctx MT L, Ctx 6d L, Ctx 45 L, Ctx 44 L, Ctx STSda L, Ctx 47l L, Ctx 8Av L, Ctx IFSp L, Ctx TE1a L, Ctx IFJa L, Ctx AVI L, Ctx FST L, Ctx FEF L, Ctx PSL L, Ctx TGd L, Ctx 55b L, Ctx 3b L, Ctx i6 8 L, Ctx p9 46v L, Ctx PHT L, Ctx PEF L, Ctx TGv L* | | | | | |
| 6 | 4048 | -13 | 10 | 6 | 7.03 | Sub-cortex |
| | CAU L, VStriatum L | | | | | |
| | *BG CAU VA L, BG CAU body L, BG BST SLEA L* | | | | | |
| 7 | 10104 | -3 | -19 | 4 | 7.03 | Sub-cortex |
| | Thal Posterior L, Thal Medial L, Thal Posterior R, Thal Medial R, Midbrain R, Thal Ventral L, Midbrain L, Thal Ventral R, Thal Intralaminar R, Midbrain, Thal Intralaminar L | | | | | |
| | *Thal PuA L, Thal MD L, Thal PuA R, Thal MD R, BStem mRt R, BStem RN R, Thal VL L, BStem mRt L, Thal VL R, BStem SC R, Thal PuM L, Thal cIL R, BStem CLi RLi, Thal MGN L, BStem STH L, Thal VPL VPM L, BStem Shen Midb Lrd, Thal CL L* | | | | | |
| 8 | 3776 | 16 | 8 | 10 | 3.35 | Sub-cortex |
| | CAU R, VStriatum R | | | | | |
| | *BG CAU body R, BG CAU VA R, BG BST SLEA R* | | | | | |
| 9 | 664 | -27 | -5 | 4 | 3.81 | Sub-cortex |
| | PUT L | | | | | |
| | *BG PUT DP L* | | | | | |
| 10 | 52256 | -1 | 42 | 42 | 3.47 | Cortex_Default_ModeB |
| | cingulate dlPFC L, cingulate dlPFC R, cingulate ACC mPFC L, cingulate ACC mPFC R | | | | | |
| | *Ctx 8BL L, Ctx 8BL R, Ctx 9a L, Ctx 9m L, Ctx SFL L, Ctx 9m R, Ctx 9p L, Ctx SFL R, Ctx 8BM L, Ctx 9a R, Ctx 8BM R, Ctx 9p R, Ctx d32 L* | | | | | |
| 11 | 16168 | -1 | -59 | 38 | 5.88 | Cortex_Default_ModeA |
| | cingulate posterior R, cingulate posterior L, parietal superior lobule L | | | | | |
| | *Ctx 7m R, Ctx 7m L, Ctx 31pd L, Ctx 31pd R, Ctx 31pv L, Ctx v23ab R, Ctx PCV L, Ctx 7Pm L, Ctx d23ab L, Ctx d23ab R* | | | | | |
| 12 | 4576 | 46 | 4 | 50 | 3.88 | Cortex_Dorsal_AttentionB |
| | somatomotor premotor R | | | | | |
| | *Ctx 55b R, Ctx FEF R, Ctx PEF R* | | | | | |

**Note.** This table is based on the results of the conjunction map with only social interaction regressors in the GLM. *t* values were calculated as the average of the *t* values in the effect maps under Text and Audio modality separately. Same applied for all tables below.

**Table S2.** Regions associated with ToM engagement under both modalities

| Cluster | Volume (mm^3) | X | Y | Z | Max t | Network |
|---|---|---|---|---|---|---|
| 1 | 462816 | -5 | -17 | 16 | 7.03 | Multiple regions |
| | visual MT+ L, auditory association cortex L, cingulate posterior R, Thal Intralaminar L, Thal Posterior L, Thal Intralaminar R, Midbrain R, Midbrain L, parietal inferior lobule L, Thal Ventral L, Thal Posterior R, cingulate dlPFC R, parietal TPOJ L, cingulate ACC mPFC L, somatomotor paracentral lobule R, cingulate vlPFC R, cingulate dlPFC L, CAU L, parietal TPOJ R, cingulate posterior L, somatomotor premotor L, Cblm vermis, parietal inferior lobule R, temporal lateral R, somatomotor premotor R, visual dorsal R, Pons L, somatomotor paracentral lobule L, Thal Medial L, Thal Ventral R, cingulate ACC mPFC R, auditory association cortex R, PUT L, cingulate vlPFC L, visual MT+ R, insula anterior L, visual dorsal L, parietal superior lobule L, PUT R, hypothalamus R, somatomotor primary L, cingulate ventral frontal L, CAU R, Cblm cortex R, Thal Medial R, hypothalamus L, Midbrain, visual ventral R, parietal superior lobule R, temporal lateral L, Cblm cortex L, GP L, insula operculum L, cingulate ventral frontal R, Thal Lateral L, Thal Anterior L, insula anterior R, visual ventral L, visual early L, VStriatum L, somatomotor operculum L, visual early R, Pons R, VStriatum R, Thal Lateral R | | | | | |
| | *Ctx MST L, Ctx MT L, Ctx STSvp L, Ctx 7m R, Thal cIL L, Thal PuA L, Thal rIL R, BStem STH R, Thal MGN L, BStem RN L, BStem RN R, BStem isRt L, BStem mRt L, BStem mRt R, Ctx PGi L, Thal VPL VPM L, Thal cIL R, Thal PuA R, Ctx 8BL R, Ctx TPOJ2 L, Ctx 8BM L, Thal rIL L, Ctx SCEF R, Ctx 45 R, Ctx 8BL L, BG CAU VA L, Thal LGN R, Ctx TPOJ2 R, Ctx 7m L, Ctx 31pd R, Ctx V4t L, Ctx SFL L, Ctx 6a L, Cblm Vermis IX, Ctx 8C L, Thal PuL L, BStem SC L, Ctx PGi R, Ctx TE1a R, Ctx 9p L, Ctx FEF R, Ctx V3CD R, BStem PTg L, Ctx 31pd L, Ctx 24dd L, Thal MD L, Ctx 6ma R, Thal VL R, Ctx STSva L, Ctx 9m L, Ctx 9m R, Ctx SFL R, Ctx 6d R, BStem VTA PBP L, Ctx STSva R, Thal VL L, Thal VA L, BStem STH L, BG PUT DA L, Ctx 9a L, Ctx 44 L, Ctx MST R, Ctx TPOJ1 L, Ctx IFJa R, Thal PuM L, Ctx AVI L, Ctx IFSp L, Ctx 8C R, Ctx 9a R, BStem isRt R, Ctx V3B L, Thal MGN R, BStem VTA PBP R, Ctx STV R, Thal PuL R, Ctx 44 R, Ctx V3CD L, Ctx FST R, Ctx 7Pm L, Ctx 9p R, Ctx 6a R, Ctx 24dv L, Ctx SCEF L, Ctx STSdp L, BG PUT DP R, hypothalamus posterior R, Ctx 45 L, Ctx 6ma L, Ctx 2 L, Ctx FST L, Thal VA R, BStem MiTg PBG L, Ctx FEF L, Ctx 47s L, Thal LGN L, Ctx 47l L, Ctx p32pr R, Ctx p32pr L, BStem SN L, BG PUT DP L, BStem SC R, Thal VPL VPM R, Ctx STSvp R, Ctx STV L, Ctx MT R, BG CAU body L, Ctx 6d L, BG CAU body R, Ctx 6v R, BStem Shen Midb Rrd, Ctx TPOJ3 R, Ctx 55b R, Ctx PFt L, Cblm IX R, Thal MD R, BG PUT DA R, Ctx AIP L, hypothalamus posterior L, BStem Shen Midb Lrd, Ctx 6mp L, BStem PAG, Ctx 8Ad R, Ctx V8 R, Ctx 7Pm R, Ctx 8Av L, Ctx LO2 R, Ctx 31pv L, BStem CLi RLi, Ctx 8BM R, Ctx PCV L, BStem MiTg PBG R, Ctx TGd R, Ctx IFJa L, Ctx TE1a L, Ctx TGd L, Ctx i6 8 L, Ctx 7PC L, Ctx V7 L, Thal CL L, Ctx p9 46v L, Cblm IX L, BG GPe L, BStem Shen Pons Lv, Ctx v23ab R, Ctx s6 8 R, Ctx FOP2 L, BStem SN R, BG CAU DA L, Ctx 47l R, Ctx LIPv L, Thal LD L, BG PUT VA L, Thal AV L, BG CAU VA R, Ctx PSL L, Ctx PEF R, Cblm CrusII R, Ctx FOP5 L, Ctx LO1 R, Ctx LO1 L, Ctx PCV R, Ctx POS2 R, Ctx 3b L, Ctx TGv L, Ctx d32 L, Ctx AVI R, Cblm Vermis VIIIb, BG PUT VP L, Ctx V8 L, Ctx V4 L, Ctx TPOJ1 R, Ctx V4t R, Ctx 8Av R, BStem Shen Pons Lcd, Ctx 24dv R, Ctx PHT L, Ctx PFt R, Ctx IFSp R, BG BST SLEA L, Cblm CrusII L, Thal PuM R, Cblm VI R, Ctx TPOJ3 L, Cblm V R, Ctx p9 46v R, Ctx 6mp R, BG CAU DA R, Ctx FOP1 L, Ctx 55b L, Ctx FOP4 L, Ctx 47s R, Ctx STSda L, Ctx 6v L, Ctx V4 R, BStem Shen Pons Rcv, Ctx d23ab L, Ctx PEF L, BG BST SLEA R, Ctx STGa R, Thal LP R, Ctx 3a L, Ctx 6r L, Ctx 4 L, Ctx 1 L, Ctx 24dd R, Ctx MIP L, Ctx IPS1 L, Thal LP L, Cblm CrusI R, Ctx TE1m L, Ctx 31pv R, Ctx d23ab R, BStem Shen Midb Rcd, Cblm VIIb R, Ctx STSdp R, Ctx V3B R, BStem Shen Midb Lc, Thal CL R, BStem PnO PnC R, BStem IC L, Ctx 23d L, Cblm Vermis X, Ctx STSda R, Ctx 46 L, Ctx p24pr L, hypothalamus anterior and tubular inferior L* | | | | | |
| 2 | 2536 | -1 | 56 | -19 | 5.20 | Cortex_Limbic |
| | cingulate ventral frontal R | | | | | |
| | *Ctx 10v R* | | | | | |
| 3 | 1904 | -49 | -25 | 22 | 3.41 | Cortex_SomatomotorB |
| | somatomotor operculum L | | | | | |
| | *Ctx OP1 L, Ctx PFcm L* | | | | | |
| 4 | 704 | 46 | -27 | 22 | 7.03 | Cortex_SomatomotorB |
| | somatomotor operculum R | | | | | |
| | *Ctx PFcm R* | | | | | |

**Note.** This table is based on the results with only ToM engagement regressors in the GLM.

**Table S3.** Regions associated with ToM demands under both modalities

| Cluster | Volume (mm^3) | X | Y | Z | Max t | Network |
|---|---|---|---|---|---|---|
| 1 | 8288 | -49 | 4 | -33 | 4.97 | Cortex_Default_ModeB |
| | temporal lateral L | | | | | |
| | *Ctx TGv L, Ctx TGd L* | | | | | |
| 2 | 6376 | 52 | 6 | -35 | 7.03 | Cortex_Limbic |
| | temporal lateral R | | | | | |
| | *Ctx TE1a R, Ctx TGd R* | | | | | |
| 3 | 5824 | -49 | 24 | -1 | 7.03 | Cortex_Default_ModeB |
| | cingulate vlPFC L, cingulate ventral frontal L | | | | | |
| | *Ctx 45 L, Ctx 47l L, Ctx 44 L* | | | | | |
| 4 | 12128 | 32 | -81 | -1 | 7.03 | Cortex_Visual_Central |
| | visual ventral R, visual MT+ R, visual dorsal R, visual early R | | | | | |
| | *Ctx VMV3 R, Ctx LO1 R, Ctx V8 R, Ctx V3CD R, Ctx V4 R, Ctx VMV2 R* | | | | | |
| 5 | 13920 | -29 | -87 | -1 | 7.03 | Cortex_Visual_Central |
| | visual MT+ L, visual ventral L, visual dorsal L, visual early L | | | | | |
| | *Ctx LO1 L, Ctx VMV3 L, Ctx V3CD L, Ctx V4 L, Ctx V8 L, Ctx MST L, Ctx LO2 L* | | | | | |
| 6 | 6408 | -53 | -31 | -5 | 6.72 | Cortex_Default_ModeB |
| | auditory association cortex L | | | | | |
| | *Ctx STSvp L, Ctx STSdp L* | | | | | |
| 7 | 520 | 44 | -65 | 2 | 5.01 | Cortex_Dorsal_AttentionA |
| | visual MT+ R | | | | | |
| | *Ctx MST R* | | | | | |
| 8 | 4768 | -51 | -59 | 30 | 4.48 | Cortex_Default_ModeB |
| | parietal inferior lobule L, parietal TPOJ L | | | | | |
| | *Ctx PGi L, Ctx STV L* | | | | | |
| 9 | 24080 | -11 | 34 | 50 | 3.36 | Cortex_Default_ModeB |
| | cingulate dlPFC L, somatomotor paracentral lobule L, cingulate ACC mPFC L | | | | | |
| | *Ctx SFL L, Ctx 8BL L, Ctx 9a L, Ctx 9p L, Ctx SCEF L, Ctx p32pr L, Ctx 9m L* | | | | | |
| 10 | 17216 | -1 | -63 | 38 | 7.03 | Cortex_Fronto_ParietalC |
| | cingulate posterior L, cingulate posterior R, visual dorsal L | | | | | |
| | *Ctx POS2 L, Ctx POS2 R, Ctx RSC L, Ctx RSC R, Ctx V6 L, Ctx DVT R* | | | | | |
| 11 | 3760 | -35 | -57 | 50 | 3.12 | Cortex_Fronto_ParietalA |
| | parietal superior lobule L, parietal inferior lobule L | | | | | |
| | *Ctx LIPd L, Ctx IP2 L* | | | | | |
| 12 | 6176 | -31 | -15 | 60 | 3.31 | Cortex_SomatomotorA |
| | somatomotor premotor L | | | | | |
| | *Ctx 6a L* | | | | | |

**Note.** This table is based on the results with only ToM demands regressors in the GLM.

**Table S4.** Regions associated with social interactions controlling for ToM under both modalities

| Cluster | Volume (mm^3) | X | Y | Z | Max t | Network |
|---|---|---|---|---|---|---|
| 1 | 3600 | 2 | -57 | -45 | 3.58 | Sub-cortex |
| | Cblm cortex R, Cblm cortex L, Cblm vermis | | | | | |
| | *Cblm IX R, Cblm IX L, Cblm Vermis VIIIb* | | | | | |
| 2 | 34976 | -7 | -81 | -27 | 3.39 | Sub-cortex |
| | visual ventral L, visual early L, Cblm cortex R, visual ventral R, Cblm cortex L, visual dorsal L | | | | | |
| | *Ctx V8 L, Ctx V4 L, Cblm CrusII R, Ctx V8 R, Ctx VMV3 L, Cblm CrusII L, Ctx V3CD L* | | | | | |
| 3 | 32264 | 58 | 4 | -15 | 7.03 | Cortex_Default_ModeB |
| | cingulate vlPFC R, auditory association cortex R, temporal lateral R, cingulate ventral frontal R, somatomotor premotor R, insula anterior R | | | | | |
| | *Ctx 45 R, Ctx STSvp R, Ctx STSva R, Ctx IFJa R, Ctx TE1a R, Ctx 47l R, Ctx IFSp R, Ctx 55b R, Ctx STGa R, Ctx STSdp R, Ctx TGd R, Ctx 44 R, Ctx 47s R, Ctx STSda R, Ctx AVI R* | | | | | |
| 4 | 27664 | -53 | -37 | 4 | 7.03 | Cortex_Default_ModeB |
| | auditory association cortex L, parietal inferior lobule L, parietal TPOJ L, temporal lateral L | | | | | |
| | *Ctx STSvp L, Ctx PGi L, Ctx STSva L, Ctx STSdp L, Ctx TPOJ2 L, Ctx TPOJ1 L, Ctx STV L, Ctx STSda L, Ctx TE1a L* | | | | | |
| 5 | 15912 | -47 | 20 | 16 | 7.03 | Cortex_Default_ModeB |
| | cingulate ventral frontal L, cingulate dlPFC L, cingulate vlPFC L, insula anterior L | | | | | |
| | *Ctx 47s L, Ctx 8C L, Ctx 45 L, Ctx 47l L, Ctx 8Av L, Ctx 44 L, Ctx IFSp L, Ctx AVI L* | | | | | |
| 6 | 1768 | -11 | 10 | 4 | 7.03 | Sub-cortex |
| | CAU L | | | | | |
| | *BG CAU VA L* | | | | | |
| 7 | 4224 | -1 | -19 | 6 | 2.56 | Sub-cortex |
| | Thal Medial L, Thal Posterior L, Midbrain R, Thal Medial R, Thal Posterior R | | | | | |
| | *Thal MD L, Thal PuA L, BStem SC R, Thal MD R, Thal PuA R* | | | | | |
| 8 | 4344 | 30 | -93 | 8 | 7.03 | Cortex_Visual_Central |
| | visual dorsal R | | | | | |
| | *Ctx V3CD R* | | | | | |
| 9 | 2032 | 12 | 10 | 10 | 4.21 | Sub-cortex |
| | CAU R | | | | | |
| | *BG CAU VA R, BG CAU body R* | | | | | |
| 10 | 41280 | -3 | 44 | 42 | 3.22 | Cortex_Default_ModeB |
| | cingulate dlPFC L, cingulate ACC mPFC L, cingulate ACC mPFC R, cingulate dlPFC R | | | | | |
| | *Ctx 9a L, Ctx 9m L, Ctx 8BL L, Ctx 9m R, Ctx SFL L, Ctx 8BL R, Ctx 9p L, Ctx SFL R, Ctx 8BM L, Ctx 9a R, Ctx 8BM R, Ctx d32 L* | | | | | |
| 11 | 9880 | -1 | -59 | 36 | 3.46 | Cortex_Default_ModeA |
| | cingulate posterior L, cingulate posterior R | | | | | |
| | *Ctx 7m L, Ctx 31pd L, Ctx 7m R, Ctx 31pv L, Ctx d23ab L, Ctx v23ab R* | | | | | |

**Note.** This table is based on the results with both social interactions and ToM engagement regressors in the GLM.

**Table S5.** Regions associated with ToM controlling for social interactions under both modalities

| Cluster | Volume (mm^3) | X | Y | Z | Max t | Network |
|---|---|---|---|---|---|---|
| 1 | 511784 | -5 | -19 | 16 | 7.03 | Multiple regions |
| | visual MT+ L, auditory association cortex L, cingulate posterior R, Thal Intralaminar L, Thal Posterior L, Thal Intralaminar R, Midbrain R, Midbrain L, parietal inferior lobule L, Thal Ventral L, parietal TPOJ L, cingulate dlPFC R, somatomotor paracentral lobule R, parietal TPOJ R, Thal Ventral R, Thal Posterior R, somatomotor premotor R, cingulate posterior L, cingulate ACC mPFC L, visual dorsal R, somatomotor premotor L, cingulate dlPFC L, visual MT+ R, parietal inferior lobule R, somatomotor paracentral lobule L, Cblm vermis, parietal superior lobule L, CAU L, cingulate vlPFC R, cingulate ACC mPFC R, PUT L, cingulate vlPFC L, Thal Medial L, visual dorsal L, temporal lateral R, somatomotor primary L, insula operculum L, insula anterior L, PUT R, Pons L, parietal superior lobule R, CAU R, somatomotor operculum L, Midbrain, cingulate ventral frontal L, Cblm cortex R, visual ventral R, Cblm cortex L, temporal lateral L, GP L, Thal Lateral L, Thal Medial R, visual ventral L, somatomotor operculum R, cingulate ventral frontal R, auditory association cortex R, visual early L, somatomotor primary R, hypothalamus R, insula operculum R, Pons R, visual early R, Thal Lateral R, VStriatum L, insula anterior R, hypothalamus L, VStriatum R, auditory early R, GP R, Thal Anterior L | | | | | |
| | *Ctx MST L, Ctx MT L, Ctx STSvp L, Ctx 7m R, Thal cIL L, Thal PuA L, Thal cIL R, BStem STH R, BStem RN L, BStem RN R, BStem isRt L, BStem mRt L, BStem mRt R, Ctx PGi L, Thal VPL VPM L, Ctx TPOJ2 L, Ctx 8BL R, Ctx 6ma R, Ctx SCEF R, Ctx TPOJ2 R, Thal VPL VPM R, Thal PuA R, Ctx 6a R, Thal rIL R, Ctx 7m L, Ctx 8BM L, Ctx V3CD R, Ctx V4t L, Ctx 6a L, Ctx 8BL L, Ctx FST R, Ctx PGi R, Ctx 31pd R, Ctx 9p R, Ctx 6d R, Ctx SFL L, Thal PuL L, Ctx 8Ad R, Ctx 24dd L, Ctx 8C L, Thal LGN R, BStem SC L, Ctx FST L, Cblm Vermis IX, Ctx 8C R, Ctx 9p L, Ctx 31pd L, Ctx FEF R, Thal MGN L, Ctx MST R, Thal VL L, BStem STH L, Ctx 7Pm L, Ctx SFL R, BG CAU VA L, Ctx IFJa R, Ctx 9m R, BStem isRt R, Ctx 9m L, BG PUT DA L, Ctx 44 L, Thal PuL R, Ctx 9a R, Thal MD L, Ctx PFt R, Ctx 24dv L, Ctx PFt L, Ctx p32pr L, Thal VL R, Ctx p32pr R, Ctx 9a L, Ctx V3CD L, Thal PuM L, Ctx TE1a R, Ctx s6 8 R, Ctx 2 L, Ctx IFSp L, Ctx 44 R, Ctx V3B L, Thal rIL L, Ctx 45 R, Ctx AIP L, BStem VTA PBP L, Ctx SCEF L, Ctx 6ma L, Ctx 6v R, Ctx FEF L, Ctx FOP2 L, Ctx AVI L, Ctx 45 L, BG PUT DP R, Thal LGN L, Ctx 24dv R, Ctx MT R, BG PUT DP L, Ctx i6 8 L, Ctx TPOJ1 L, Ctx 6mp L, BStem PTg L, Ctx LO2 R, Ctx POS2 R, Ctx 8BM R, Ctx 7Pm R, Ctx STV R, BG CAU body R, Ctx 6d L, Ctx 7PC L, Ctx 6mp R, BStem VTA PBP R, Ctx OP1 L, BStem PAG, Ctx 47l L, Cblm IX R, Ctx STV L, Ctx FOP1 L, BG CAU body L, Ctx PEF R, Ctx LIPv L, Ctx 6r L, Thal VA L, Ctx 8Av L, Ctx TPOJ3 R, Ctx IFJa L, Ctx STSdp L, BStem CLi RLi, Ctx 7PL L, Ctx p9 46v L, Ctx PCV L, Ctx V4t R, Ctx 31pv L, BStem Shen Midb Rrd, Ctx 47s L, Thal MGN R, Ctx 8Av R, Ctx 55b R, Ctx V8 R, Cblm IX L, Thal VA R, Ctx IPS1 L, Ctx STSva L, Ctx v23ab R, BStem Shen Pons Lv, BStem Shen Midb Lrd, Ctx TGd L, Ctx V7 L, Ctx 3b L, Ctx PCV R, BG GPe L, Ctx LIPv R, Ctx TGd R, Cblm Vermis VIIIb, BStem SC R, BStem SN L, Ctx POS2 L, Thal LD L, BG CAU DA L, BG PUT DA R, Thal MD R, Ctx LO1 R, Ctx V8 L, Ctx TE1a L, Ctx p9 46v R, Ctx LO1 L, Ctx PFcm R, Cblm CrusII R, Ctx PSL L, BStem MiTg PBG R, Cblm V R, BG PUT VP L, Ctx FOP4 L, Ctx FOP5 L, Ctx PFcm L, Ctx 47l R, Ctx PEF L, BG CAU VA R, BStem Shen Pons Lcd, Ctx TGv L, Ctx d32 L, Ctx PHT L, Ctx STSva R, Thal PuM R, BG PUT VA L, Ctx MIP L, BG CAU DA R, Ctx AIP R, Cblm VI R, Ctx V4 L, Ctx 24dd R, BStem MiTg PBG L, BStem SN R, Ctx 6v L, Cblm CrusII L, Ctx 2 R, Ctx 55b L, Ctx 7PL R, Ctx VIP L, hypothalamus posterior R, Ctx V3B R, Ctx 4 L, Ctx FOP2 R, BStem Shen Pons Rcv, Ctx d23ab L, Ctx V4 R, Ctx 46 L, Ctx p24pr L, Ctx IFSp R, Ctx 1 L, Thal LP R, Cblm VIIIa R, Ctx IP0 L, Ctx 31pv R, Ctx 3a L, BG BST SLEA L, Thal CL L, Ctx POS1 R, Ctx AVI R, Ctx TPOJ3 L, hypothalamus posterior L, Cblm VIIb R, Ctx d32 R, Cblm CrusI R, BG BST SLEA R, Ctx RI R, BStem PnO PnC R, Ctx TE1m L, Ctx i6 8 R, Ctx 47s R, Ctx STSvp R, Ctx TPOJ1 R, Ctx d23ab R, Thal LP L, Cblm Vermis X, Ctx a32pr L, Ctx FFC R, Ctx 9 46d L, BG GPe R, Thal CL R, Ctx s6 8 L, Ctx 5mv L, Ctx 23d L, Cblm X R, BStem Shen Pons Rcd, BStem Shen Midb Lc, Thal AV L, Ctx LO2 L, Ctx PFm L, BStem Shen Midb Rcd* | | | | | |

**Note.** This table is based on the results with both social interactions and ToM engagement regressors in the GLM.

#### Texts of all narratives used in the study

In the texts below, each paragraph contains a narrative part that was constructed as one of the 36 dramatic situations (Polti, 1917) and presented in one experimental trial.

##### Narrative #1

One day at work, Margaret walked up to her partner, Al, to ask him to talk to the chief of staff at the local hospital where she worked as a nurse.

While waiting outside the chief of staff's office, Al heard nondescript shouting and yelling, there was a loud clatter in the room and then an eerie silence. An unfamiliar employee ran hastily out of the office with a terrified look on their

face.
Al explained to the chief of staff that Margaret wanted to be excused from her absences last week. The chief of staff, Susan, revealed that she wanted to fire Margaret and never see her again. Susan and Al had been having an affair and she wanted Margaret “out of the picture”. Al was conflicted but agreed it would be for the best.
Weeks passed and nobody had seen Margaret. Coworkers and friends searched the nearby city looking for her. One day on patrol in a nearby town, a police officer heard someone crying for help in the basement of a house and went in to investigate. Margaret yelled that she was being held captive and needed help.
In a rush, the police officer broke down the door and was attacked by a man in the house. Without hesitation, the officer shot the man and the man fell to the floor.
The officer rushed to the basement, untied Margaret, and took her to safety. After being notified of the situation, Al rushed to meet Margaret as to not raise any suspicion. When he arrived, he saw Margaret pointing and yelling at Susan who had been handcuffed and was talking to the police. Shortly after, the police took Susan away in a police car. Al was devastated to see her being taken away, as Margaret sneered with disdain.
Weeks later, Al went to county lockup to speak with Susan. After briefly catching up, he asked Susan about what happened at the house. Without realizing that their conversation was being recorded, Al foolishly mentioned their plans to get rid of Margaret. He even went as far to ask about what to do next. Within days, the two of them ended up confessing about their plot.
Shortly after, Margaret heard about Al’s confession and infidelity on the local news. She was furious that she did not know what was going on and hurt that Al had been acting so selfishly behind her back.
Margaret went to a local bar to drown out her pain. After a few drinks at a quiet bar, she went to the back of the bar to use the facilities. On her way back, she heard someone approaching her from behind with a shuffled gait. She turned around and realized it was the man that was keeping her captive! Before she could make a sound, he covered her mouth with a rag and pulled her out the back of the bar – never to be seen again.

#### Narrative #2

Linda and Amy had been best friends since kindergarten, they even found homes on the same street in the forest on the edge of town. Their biggest fight had been over who gets the bigger slice of cake, until Amy married Dan. Linda loathed Dan, she thought he was manipulative and had a violent temper. She only put up with him because Amy loved him.
This feeling of hate between Amy and Dan was mutual, Dan didn’t like how much time his wife spent with her and thought Amy was creating distance between their marriage. That’s why Dan decided to try and tarnish Linda’s reputation with his wife, in hopes it would get rid of her. He came home one afternoon and exclaimed that Linda had tried to kiss him. Amy was taken aback and immediately called Linda to confront her about these accusations. When Linda arrived, she denied having made a move on Dan, but discovered Dan had enlisted the help of his friend Will who said he “witnessed” it all happen.
Since it was their word against her own, she decided to go home and admit defeat. She knew how much it would hurt Amy to know the truth.
Months later Dan had taken up a mistress, he was growing bored with his marriage ever since he got rid of Linda and was only still with Amy for her family’s money. In a hotel room in a nearby town, Dan and his mistress, Nicole, conspired to secretly kill Amy while still obtaining her fortune.
They decided to have Dan invite her to the hotel room under the guise of a romantic date. Once she arrived, Nicole was there to tie her up. Once she was secured to a chair, Nicole demanded Amy's bank passwords. Amy obliged and gave her access to her bank accounts.
Nicole transferred all of Amy’s money into Dan’s bank account despite Amy’s pleading for her to stop.
Once successful, the mistress knocked Amy out with a forceful blow to the head. After seeing Amy’s body lying limp on the ground, she was suddenly filled with regret over her actions. Nicole quickly rushed her to the hospital, realizing she had only been jealous of Amy’s marriage with Dan and that she didn’t deserve to die.
Once Amy awoke from consciousness at the hospital, she was met by Linda. Linda rushed over as soon as she was called, finding out she was still Amy’s emergency contact. Nicole apologized and explained to Amy and Linda what had happened at the hotel. Amy was taken aback when she realized her husband's betrayal.

She decided it was best to let Dan run off with her money if that meant he was out of her life forever. Amy and Linda continued to be best friends, and years later she went on to marry a kind man with a loving family.

#### Narrative #3

It was just another Sunday afternoon for Lucy, she was on her way to the farmer's market to pick up some produce. When she arrived at the old train depot that housed the market, she noticed two men fighting on the steps. Before she could call for help, one of the men crumpled to his knees. The other man dropped an object that appeared to be a knife and ran off. When Lucy approached the crime scene, she realized that the man who had been murdered was the Mayor and that he had a gunshot wound.

On the ground beside him was a pen that she had mistaken for a knife. Lucy looked for the gunman and saw a suspicious man in black with distinct pointy features darting into an alley across the street. Cops who were nearby chased the man who had fled and arrested him.

But Lucy knew the charged man was innocent, which meant the killer was free. Lucy vowed to find out who the real murderer was and to stop him at all costs.

The next day the mayor's successor, Mr. Smith, a man who was very unpopular due to his strong ties to the oil companies, took office. After a long day of brainstorming about the mysterious killer, Lucy decided to attend his welcome ceremony in the city. During the mayor's speech, she noticed one of the security guards looked strangely familiar. It dawned on her that this new mayor had motive to kill and that the man in black could've been the guard working for him.

Lucy decided to confront the mayor after his ceremony. But when she tried to accuse him, the security guards threatened to hurt her if she went to the police with her theory. Before she could say another word, a man pulled her from the crowd.

The man in question was another security guard of Mayor Smith's, who took Lucy into a hidden alleyway. His name was Max and he quickly explained that he was a part of the initial plot to kill the former Mayor, but he regretted his involvement.

He told Lucy about the Mayor's secret hideout in the forest that was used as a base for the Mayor's plots. They formed a plan to infiltrate the base with Lucy's brother Alex, a cop for the city, so they could find evidence of his crimes and arrest Mayor Smith. When the three arrived at the hideout, Max suggested they enter through the back. Once they snapped the lock off and opened the door, men grabbed hold of Lucy and Alex. The bodyguard in black came out and ordered the men to kill the cop immediately so he wouldn't talk. The man holding him instantly shot him in the chest.

Lucy pleaded with the guard to let them go so she could take her brother to the hospital, she said she would give up her mission against the mayor.

The guards laughed at her request and Lucy suddenly realized that they must have been tipped off on when they'd be at the hideout. She turned to Max and accused him of betraying them. Max shrugged and confessed to his disloyalty, he joined the other bodyguards in line and exclaimed neither of them were leaving this hideout alive. Resigned to her fate, Lucy closed her eyes and allowed the guards to kill her and any hope the former Mayor had of retribution.

#### Narrative #4

Luke always thought of himself as an honest man. That's why during his evening jog around the swamp, he pondered his recent affair with the receptionist at work. Luke looked up at the sky and asked God why he was so weak and unfaithful. He thought of the impact this would have on his family and told God of his conflicted feelings toward telling his wife Marta.

Suddenly, Luke heard a woman call out his name and when he turned around, he saw Carly, his receptionist, across the road with a gun in her hand. Carly ran to him and explained how she had followed him here to confess her love. She exclaimed if Luke didn't love her back that she would kill him and his family. Carly explained that the only way they could be together is to kill Marta.

Luke thought of his child and felt like this was his only option. He agreed to the plan and called his wife to meet him at the swamp to watch the sunset with him. When she arrived, Marta called out for her husband, but found Carly waiting for her. She immediately drew her gun and shot her in the chest, knocking her unconscious.

Luke ran to Marta and filled with regret over his decision to go along with Carly's plan, picked her up and rushed her to the hospital. The doctors tried to help Marta by taking her into emergency surgery but failed to save her life. Once the doctor called the time of death, Luke fell to his knees and sobbed over his rash decision.

He then saw Marta's brother Zach enter the hospital. Zach ran over to Luke and punched him. Zach screamed at Luke that it's his fault his sister is dead and that he knew about the affair. Luke quickly tried to explain that he felt horrible about Marta's death and that Carly had manipulated him into going along with her plan. Zach, still angry at his brother-in-law, realized his mistake in judgement and shifted his anger toward Carly.

After the tension with Zach cooled down, Luke thought of when he should call his son and inform him of his mother's passing. He would be out of school any minute and should hear the news from him. Luke reflected on his decision and realized it was necessary to sacrifice his wife to ensure their child's safety, and that Marta had told him to above all else to protect him.

The police showed up at Carly's door a few days later, having matched the prints found on the murder weapon to hers. She was charged with first degree murder and life in prison. Years passed and she still thought of her love, Luke, and how she missed him. Until one day he showed up during visiting hours.

Luke explained he needed closure and told her how much he hated her for killing his wife. Carly was heartbroken over this because she had held on to this delusion that Luke still loved her after all these years. Once back in her cell she began to plot how to get back at Luke, she blamed him for her lifelong sentence and vowed to break out. She wanted to make him feel just as miserable as she did, and she knew the only way to do that was to take what he loved most, his son. She flirted with a guard who she knew was sweet on her and convinced him to help her escape.

Once outside the prison walls, she went to the school Luke's son attended and fooled the faculty into thinking she was his aunt. She then took the child and fled with him, never to be seen again.

#### Narrative #5

William and Johnny are brothers-in-law who had been cellmates for years. Both were facing the death penalty in the coming week. They had spent the last year planning a jailbreak together to avoid their looming death.

When the day came to make their escape, everything was going according to plan. They had successfully stolen the keys from the guard and had made their run toward the exit. As they approached the final gate, William looked over his shoulder and saw the guards catching up. He thought of his own freedom and being reunited with his wife and child. Just as the guards were about to capture them, William tripped Johnny and he fell to the floor. This distraction gave William just enough time to reach the exit and escape.

When William got to the street and met his wife, Shannon, with the car, she asked where her brother was. Seeing her husband's facial expression become grim, she quickly discovered William had left him behind.

Once the two returned home, Shannon decided to take a walk in the park next to their house to process the betrayal by her husband. She met up with her friend Robert, who had been helping her take care of her child and comfort her while William had been in prison. She shared her grievances with Robert as he hugged her and in a moment of weakness the two kissed.

When Shannon returned home, she discovered William had taken their child. He left a note explaining he had sacrificed Johnny, her brother, so he could provide their child with a father. He said that she would never understand so he had to leave.

Robert and Shannon quickly made a plan to get the child back from William.

They found out William had met up with his old criminal partner Aaron, who agreed to let William and their son stay at his beach house. Shannon remembered Aaron and knew William still in debt to Aaron financially; trying to repay Aaron is what initially landed William in jail. Shannon knew she had to get there quick before Aaron did anything to William, or their child.

Robert knew he needed to help Shannon and was willing to do what he knew Shannon couldn't, kill William, if it meant saving the child.

When they arrived at Aaron's beach house, William saw them immediately. He made a run for it with the child, and both Shannon and Robert quickly caught up to them. When the kid realized it was Robert and Shannon who had been chasing them, he immediately ran away from William and into Robert's arms. William saw this and was heartbroken, wishing he had not missed so much of his child's life. He could see how happy his child was to see Robert. Aaron suddenly arrived and attempted to take the child from Robert. William lunged at him and yelled at Robert and Shannon to run away with the kid. William knew his child would be better off with them, so he stayed back to buy them time to escape.

#### Narrative #6

James's and Nancy's son Peter had been kidnapped recently. James told Nancy to stay at home and that he would search the whole town for the son, no matter how long it took. For days James had been searching all through town to find Peter, calling his name and hoping to hear his son's voice call back. One day while searching through an abandoned building, he finally heard his son yelling to him. He followed the voice and found his son tied to a chair. After looking around to make sure no one else was there, he ran to his son, untied him and led him out of the building into the main part of town. While running through town on their way home, James and Peter ran into James's good friend Matthew. Matthew was so happy that James had found Peter, but he warned them that the kidnappers would soon see that Peter was missing and they would come for James and Peter this time. Matthew suggested that they hide out at his property in the city far from town.

James was furious that the kidnappers had taken his son and that they now had to flee and hide. He told Matthew to take Peter because he was going to find the kidnappers and take revenge on his own.

When James returned to their home, he saw his wife standing by the swamp near the house with two other men. Before he could react quick enough, he saw one of the men strike his wife and she fell to the ground.

James ran to Nancy's side and knelt before her crying while the other men were standing above him feeling no remorse.

Suddenly, one of the men grabbed Nancy and despite the fact that James fought him, the man was able to overpower James and took Nancy and threw her body into the swamp.

James made his way to Matthew's house in the city to meet back up with his son. When he arrived, he told Matthew about Nancy's death. Matthew explained that he was sorry, but he thought that Peter should stay with Matthew and his wife because he needed a mother and father. In his state of sorrow and depression, he agreed that he did not have the capacity to take care of Peter and that it was best for him to stay with Matthew and his wife.

After the events of the last week or so, James felt he had suffered everything someone could.

Over the next couple weeks, the tragedies James had endured began to drive him mad and he spent the rest of his life alone and in agony and pain.

#### Narrative #7

Anna and her husband, Glenn, were in prison for robbing a convenience store. During their time in prison, Anna met Jeff, a man who was also convicted of robbery. The two of them developed a strong relationship, which turned into a romantic affair; Glenn was not aware of Anna's infidelity. One day Glenn saw Anna walking down a hallway by herself, so he followed her as she made her way to a storage closet. After she entered the closet, Glenn waited ten minutes and then went and knocked on the door. After a minute had gone by, Anna opened the door and was surprised to see Glenn standing before her. He pulled her out of the room and saw that her shirt was inside out and backwards. He grabbed her by the arms, started shaking her, and asked her what she was doing in the closet. His shaking became so violent that he shoved her to the ground. As he was about to hit her, Jeff came out of the closet and grabbed Glenn's hand before he could touch Anna again.

Jeff kneeled by Anna's side and took her hand. They discussed that something had to be done about Glenn.

Glenn could not believe what he had done and ran from them in a fit. Jeff and Anna avoided Glenn for the remainder of their sentences.

Six months later, all three of them were out of prison. Alone in the park, Glenn spoke to God and contemplated how he would atone for his sins and get his wife back.

Just then, he saw Anna walking by herself through the park; he had found her, and this was his opportunity to win her back.

Glenn ran to Anna and explained that he still loved her and had no intention to hurt her, but he wanted her back and would do anything to have her back in his life. Anna clutched her stomach and screamed that she was in pain. Glenn had no idea what was happening but swept her up and carried her to the hospital.

At the hospital, Glenn and Anna were met by Jeff, who had learned of her condition. The doctor explained that Anna was pregnant.
Anna and Jeff wanted to keep the baby but Glenn, who still wanted to be with Anna said that if the baby was Jeff's that she should terminate the pregnancy and be with him.
Anna turned to Glenn and told him that she didn't love him anymore, she loved Jeff, and that she and Jeff were going to keep the baby.

#### Narrative #8

Gloria and her brother, Jim, had been planning a month-long vacation together to a small town in Europe. The day before their trip, Gloria informed Jim that she couldn't go on the trip because her husband, Steve, no longer wanted her to go.
Jim understood but decided to go on the trip without her. When Jim did not return after a month, Gloria found out that Jim had been abducted.
Gloria was devastated and looked to Steve for support. Steve was not supportive in the way that she needed, so she turned to a coworker named Derek and he was there for her in every way possible. The two of them started spending a lot of time together and quickly developed a romantic relationship. Gloria and Derek decided that they wanted to be together and that Steve was in the way.
One day, Gloria and Derek were having dinner in the city together. Steve found them and confronted them about the affair. Steve pled for Gloria to stay with him, but she refused and demanded a divorce.
Steve turned to Derek and begged him not to take his wife from him; but Derek refused because he was in love with Gloria.
Gloria exclaimed that it was her decision and that she was going to choose Derek over Steve.
The next day, Gloria and Derek fled to Gloria's family's beach house to be alone. When they got to the beach house, they found Gloria's brother, Jim, tied up to a chair.
Apparently, he had never left for Europe but was being held captive. Gloria rushed to him, but as she was about to untie him, Jim's abductor seized Gloria and threw her to the ground.
In defense of Gloria and Jim, Derek grabbed a lamp and hit the abductor so hard that he died. Gloria released her brother and the two were finally reunited.
